# Collagen Rope Trick: Cell-Laden Fibre Assembly at Liquid Interfaces

**DOI:** 10.64898/2026.05.20.726709

**Authors:** Asuka Yamada, Koichi Hattori, Ayumi Watanabe, Yucheng Shang, Andrij Pich, Shiro Kitano, Michiya Matsusaki

**Affiliations:** TOPPAN HOLDINGS INC. Business Development Division, Technical Research Institute, Takanodaiminami, Sugito-machi, Saitama 345-8508, Japan; Joint Research Laboratory for Social Implementation of Cultured Meat, Yamadaoka, Suita, Osaka 565-0871, Japan; Division of Applied Chemistry, Graduate School of Engineering, The University of Osaka, Yamadaoka, Suita, Osaka 565-0871, Japan; Joint Research Laboratory (TOPPAN) for Advanced Cell Regulatory Chemistry, Yamadaoka, Suita, Osaka 565-0871, Japan; Institute of Technical and Macromolecular Chemistry, RWTH Aachen University, Worringerweg 2, D-52074 Aachen, Germany; DWI-Leibniz-Institute for Interactive Materials, Forckenbeckstraße 50, D-52056 Aachen, Germany

## Abstract

Tissues and organs in living organisms represent centimeter-scale hierarchical architectures comprising nano-to microscale, uniaxially aligned extracellular matrix (ECM) fibres with high mechanical strength, integrated with cellular components, as exemplified in tendon, skin, cartilage, bone, and blood vessels^1^. Here, we present a liquid–liquid interfacial spinning method to produce highly uniaxially aligned, centimeter-scale collagen fibres. The dried fibres exhibit exceptional mechanical properties, with fracture strength of 280 MPa, Young’s modulus of 6 GPa, and toughness of 17 MJ m⁻³, comparable to spider silk and tendon collagen, and exceeding supramolecular and double-network hydrogels^1^. Incorporating living cells into the collagen solution yielded centimeter-scale, cell-laden aligned fibres, with densely adherent, uniaxially aligned cells and over 80% viability. Myoblast-laden fibres recapitulate biological features of fibrotic muscle tissues, as observed in type II diabetes^2^. Interfacial collagen assembly further enables fabrication of dimension-controlled constructs, like 2D sheets, 0D capsules, and 1D tubes, thus providing modular building blocks for centimeter-scale 3D tissues and organ-like structures. This approach offers a versatile platform to engineer mechanically robust, cell-laden tissues with controlled hierarchical architecture.

---

Fibrous tissues in mammals—including skeletal muscle, cardiac muscle, tendons, and nerves—exhibit hierarchical, centimeter-scale architectures composed of micrometer-scale, anisotropically aligned extracellular matrix (ECM) fibres and cells^3^. Replicating these structures in vitro remains a longstanding challenge in bioengineering. An ideal scaffold should support diverse cell types, maintain stable cell–ECM interactions, and promote topographical alignment of adherent cells under cytocompatible conditions^4,5^. Recently developed approaches include melt electrowriting of fibre bundles^6^, twisting cell sheets^7^, chemically crosslinking recombinant elastin^8^, microfabrication techniques using microfluidics^9^ and filamented light patterning^10^ enabled the formation of cell-laden constructs. However, these strategies remain limited in scalability, precise cell–ECM alignment, cytocompatibility, and mechanical robustness, and cells typically attach only to material surfaces. These challenges highlight the need for new strategies to engineer hierarchical fibrous scaffolds that faithfully mimic native tissue architecture and functionality.

We recently developed a simple method to fabricate cell-laden viscous matrices by mixing collagen, heparin, and cells in Tris-HCl buffer (pH 7.4) for just 1 min, enabling the formation of three-dimensional (3D) cancer–stromal or hepatocyte tissue constructs^11,12^. Microscope observation of the resulting matrix in the absence of cells revealed micro-to millimeter-scale random fibrous structures (Extended Data Fig. 1a). Based on this observation, we hypothesized that nanometer-scale collagen assembly could be accelerated through interactions with polyanions, as heparin is a typical strong polyelectrolyte. To test this, we optimized collagen and heparin concentrations, the composition of polyanion components, and the contact conditions between collagen and polyanions.

Here we report a liquid–liquid interfacial spinning method to generate cell-laden collagen fibres, analogous to the “nylon rope trick”^13^, an interfacial polymerization of 6,6-nylon. Unlike nylon, however, interfacial spinning of proteins—particularly ECM with embedded cells—has not been realized. Remarkably, collagen molecules rapidly accumulated at the interface between collagen in acetic acid and poly(acrylic acid) (PAA) in phosphate-buffered saline (PBS), forming stable films that could be pulled into the air even under cell-containing conditions. This phenomenon enables the fabrication of dimensionally controlled tissue constructs, which can serve as modular building blocks for larger tissues or organ-like structures.

## Interfacial Spinning and Structural Characterization

Dropping a collagen solution in acetic acid (pH 4) onto PAA in PBS (pH 4) immediately generated an opaque interfacial film (Fig. 1a and Supplementary Video 1). This film was sufficiently robust to be continuously drawn into air, enabling fibre spinning over lengths of approximately 15 cm. Phase-contrast microscope observations suggest that the film formation occurred within 500 ms (Supplementary Video 2). Among 15 polymers screened, including neutral polymers, cationic and anionic polyelectrolytes, PAA exhibited the highest capacity for collagen fibre formation (Extended Data Fig. 1a,b). Since collagen molecules carry a weak positive charge at pH 3–4, electrostatic interactions likely drive collagen assembly at the liquid–liquid interface, as PAA molecules carry a weak negative charge under the same conditions (Fig. 1a, inset). This interfacial assembly and film formation strongly depends on the concentration and pH balance between collagen and PAA (Extended Data Fig. 1c-e). Continuous vertical drawing of the optimized interfacial film enabled instantaneous spinning of uniform, macroscopic fibres up to 15 cm in length (Fig. 1a, Extended Data Fig. 2a and Supplementary Videos 3,4). A highly aligned, tendon-like hierarchical architecture^14^-comprising microfibrils (50–500 nm in diameter), fibrils formed from bundles of microfibrils (1–10 µm), and larger bundles (>10 µm) assembled from fibrils-was observed in both dry and wet states (Fig. 1b) using phase-contrast (Ph), atomic force microscopy (AFM), and scanning electron microscopy (SEM), respectively. SEM images of lyophilized native collagen prior to use indicate that the spinning process generated a uniaxially aligned morphology (Extended Data Fig. 2b), while pitch measurements across hierarchical levels further corroborate this precise topographical alignment. (Extended Data Fig. 2c, d).

**Fig. 1.**
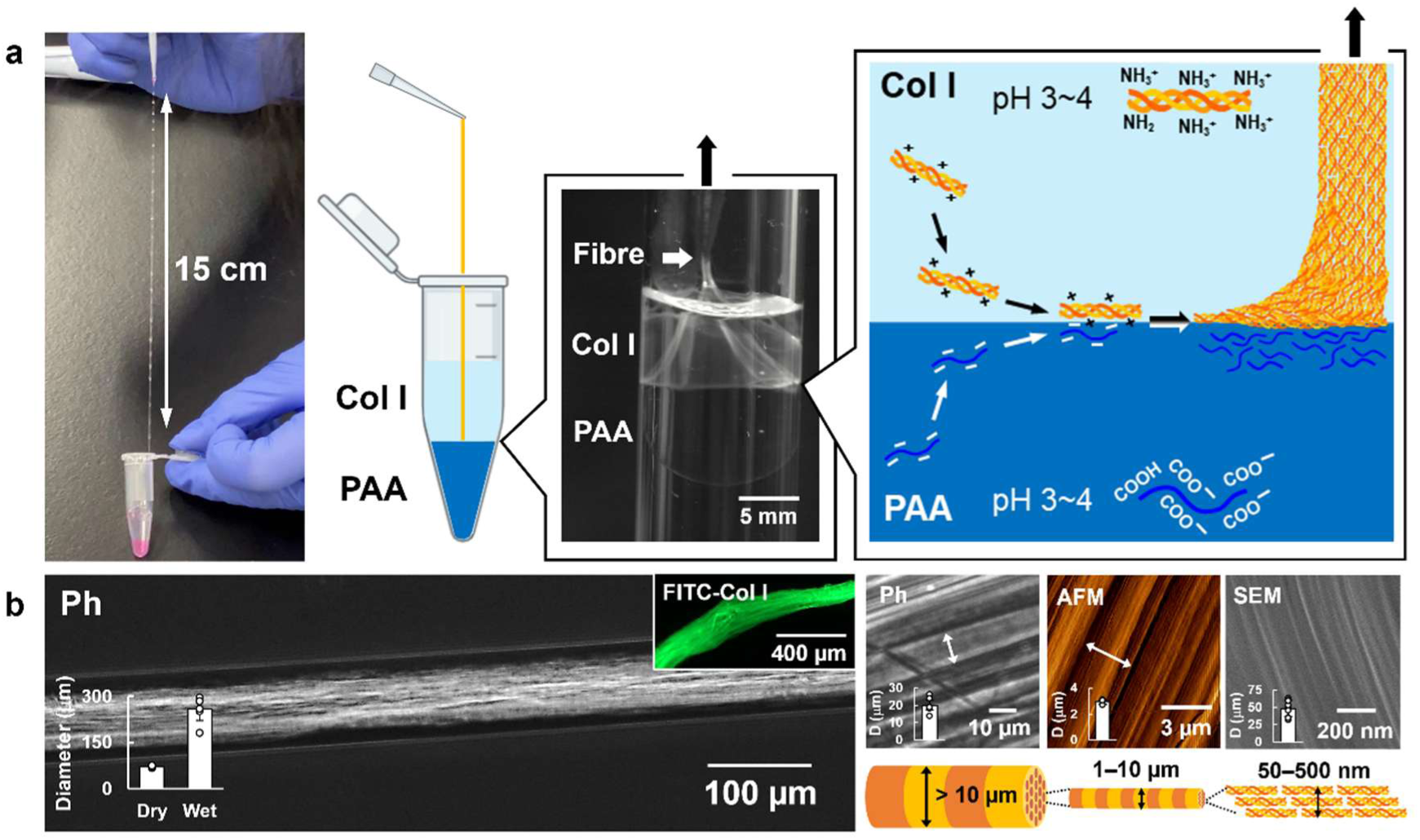
Interfacial spinning process of collagen fibres. a, Photograph (left), schematic (center), and high-speed camera snapshot (1,000 fps) (right) of the collagen fibre spinning process over 15 cm at the interface between Col I in 5 mM acetic acid solution (pH 4) and PAA in PBS (pH 4). The schematic illustrates the proposed mechanism of collagen assembly: negatively charged PAA molecules (COO⁻) interact electrostatically with positively charged collagen molecules (NH₃⁺) near the interface. The associated collagen molecules assemble into microfibrils/bundles through dehydration during spinning under ambient air conditions. **b,** Phase-contrast (Ph), atomic force microscopy (AFM), and scanning electron microscopy (SEM) images of dried collagen fibres obtained by overnight vacuum-drying. Fluorescence microscopy images of FITC-labeled collagen fibres after washing with ultrapure water are shown in the inset. Fibre diameters measured from Ph, AFM, and SEM images are shown for samples dried (Dry) or not dried (Wet) under vacuum overnight (*n* = 3–6).

To verify that this hierarchical assembly preserves the native molecular conformation of collagen, we performed spectroscopic and thermal analyses. X-ray diffraction (XRD) patterns of the fibres exhibited a characteristic equatorial reflection at 2θ ≈ 7–8°, corresponding to a lateral packing distance of ∼1.1 nm, consistent with native collagen^15^. This indicates that the highly ordered lateral assembly of the triple-helical molecules was maintained throughout the spinning process (Extended Data Fig. 2e).

Fourier-transform infrared (FT-IR) spectroscopy revealed characteristic amide bands consistent with native collagen, while PAA-derived peaks were barely detectable (Extended Data Fig. 2f). Thermogravimetric analysis (TGA) further supported this, with *T*_10_ and *T*_50_ values nearly identical to those of collagen, indicating that collagen is the primary component of the fibre (Extended Data Fig. 2g). This was visually confirmed by the selective retention of FITC-labeled collagen after extensive washing (Supplementary Fig. 1). Notably, these structural hallmarks translate into exceptional mechanical properties. Dynamic rheological measurements demonstrated a storage modulus (*G′*) several orders of magnitude higher than that of conventional physically crosslinked bulk collagen gels (Extended Data Fig. 2h). Tensile testing further revealed fracture strength, Young’s modulus, and toughness comparable to - or exceeding - those of existing natural and synthetic fibres (Fig. 2a; Extended Data Fig. 2i–m)^16,17^. These results demonstrate that dense packing driven by electrostatic interactions confers exceptional structural stability, without the need for chemical crosslinkers.

**Fig. 2.**
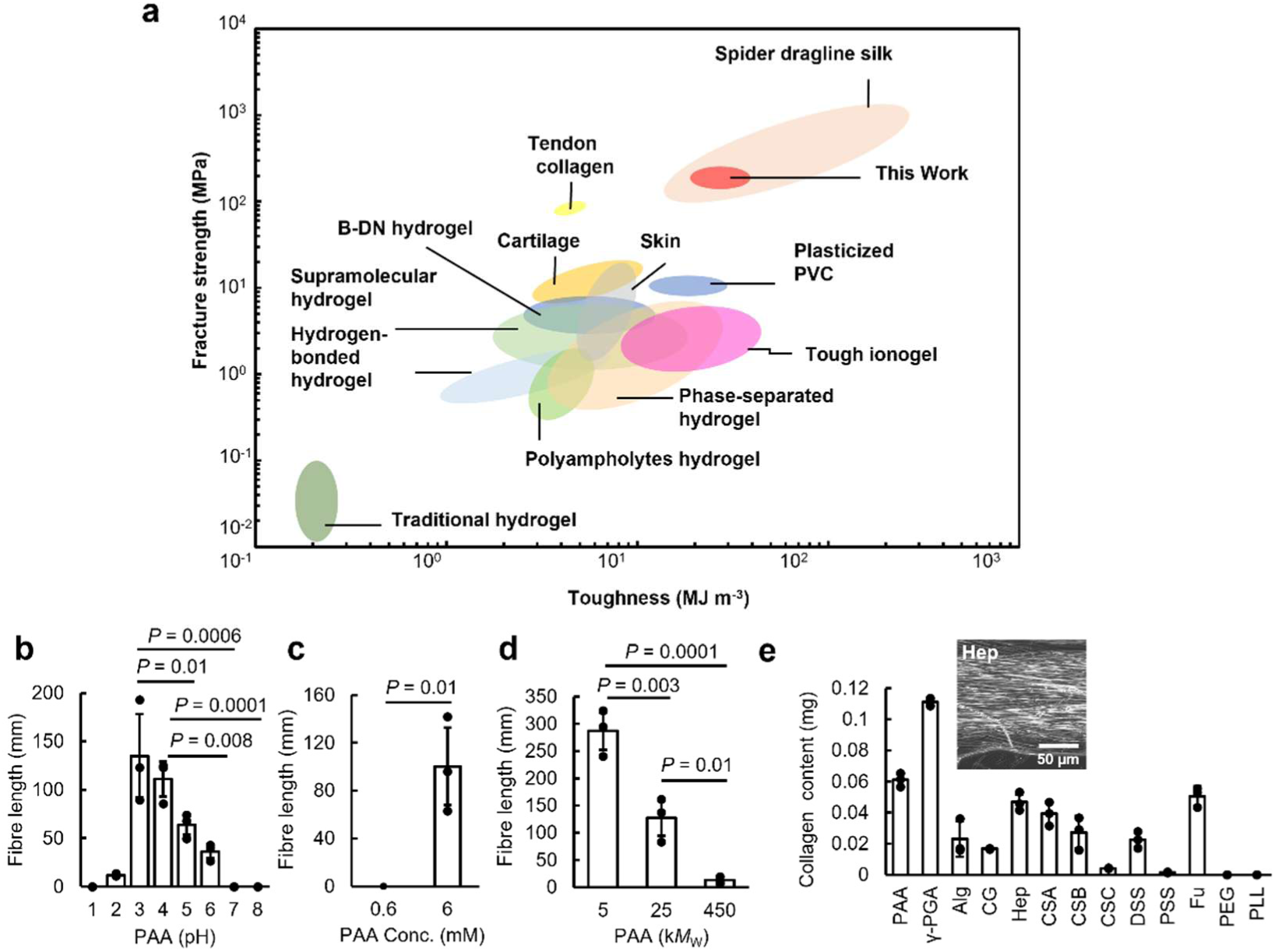
Variation of collagen fibre properties. **a** Comparison of this work (red area) with various gels and fibres, plasticized polymers, and biological tissues—including skin, cartilage, tendon collagen, and spider dragline silk—in terms of fracture strength and toughness using an Ashby-type plot. Reference material data were adapted from Wang et al.^1^, with individual data points verified and refined from the literature (Supplementary Table 1). **b–d,** Fibre length plotted against pH (b), concentration (c) and molecular weight (d) of PAA solution (*n* = 3). Statistical significance was determined by one-way ANOVA with Tukey’s post hoc test. **e,** Collagen content measured by Sirius Red staining using different counter polymer components in PBS (*n* = 3). Accumulated collagen at the interface was corrected, washed with ultrapure water, and vacuum-dried overnight. γ-PGA, poly(*γ*-glutamic acid); Alg, alginic acid; CG, carrageenan; Hep, heparin; CSA–CSC, chondroitin sulfate A–C; DSS, dextran sulfate; PSS, poly(styrenesulfonate); Fu, fucoidan; PEG, poly(ethylene glycol); PLL, poly(_L_-lysine).

### Mechanism of Interfacial Assembly and Material Universality

Next, we identified the physicochemical factors governing this interfacial spinning and elucidated the underlying mechanism. Systematic evaluation of PAA solution parameters, like pH, concentration, and molecular weight, revealed that fibre formation was maximized under acidic conditions (pH 3–4), particularly with high-concentration of low-molecular-weight PAA (Fig. 2b–d). This pH dependence strongly correlated with the amount of collagen accumulated at the interface across a broad pH range (1–11) (Extended Data Fig. 3a), as well as with the ionization states of carboxyl groups (COO⁻) on PAA and amino groups (NH₃⁺) on collagen, suggesting that an appropriate charge balance is essential for complex formation (Extended Data Fig. 3b). To assess the role of electrostatic interactions in this process, we quantified collagen accumulation at the interface using 15 selected polymers with different chemical structures, including different anionic polyelectrolytes, neutral poly(ethylene glycol) (PEG) and cationic polyelectrolyte poly(_L_-lysine) (PLL) (Fig. 2e, Extended Data Fig. 3c). Polyanions with high charge densities, such as PAA, poly(γ-glutamic acid) (γ-PGA), heparin, chondroitin sulfate A and B (CSA and CSB), dextran sulfate sodium (DSS), and fucoidan (Fu), induced pronounced collagen condensation at the interface. In contrast, polyelectrolytes with low charge densities, as well as neutral (PEG) or cationic (PLL) polymers, showed minimal collagen condensation. Thus, electrostatic condensation at the liquid–liquid interface is the dominant driving force of this process. However, fibre length does not scale with interfacial accumulation; PAA, in particular, yields exceptionally long fibres (Extended Data Fig. 3d). Although more electronegative polymers, such as dextran sulfate (DSS) and heparin, form complexes more rapidly than PAA, electrostatic repulsion arising from excess anionic charges likely extends the polymer chains and rigidifies the interface, thereby increasing resistance to drawing and promoting frequent rupture^18^. Furthermore, highly branched polymers such as Fu show high interfacial accumulation but low spinnability, likely owing to slow interface formation kinetics and lack of entanglements^19^. PEG fails to form an interface altogether (Extended Data Fig. 3e). In contrast, linear, flexible PAA achieves an optimal balance between rapid interfacial formation and continuous drawability of collagen fibres.

Another notable feature of this approach is its generality. Robust long-fibre formation is achieved not only with bovine skin–derived type I collagen, but also with cartilage-derived type II, skin-derived type III, as well as fish-derived collagen and atelocollagen (Extended Data Fig. 3f). All systems exhibit densely packed, uniaxially aligned morphologies (Extended Data Fig. 3g). Furthermore, extending the analysis to other proteins interacting with PAA shows that bovine serum albumin (BSA), laminin and fibronectin do not form complexes, whereas gelatin exhibits pronounced aggregation (Extended Data Fig. 3h). These results indicate that the interfacial behavior is governed not by specific three-dimensional conformations or amino acid sequences, but by intrinsic charge characteristics, notably cationic charge density of collagen and gelatin. This supports the broad applicability of our platform for the fabrication of diverse engineered tissues using tissue-specific extracellular matrices.

### Cell-laden Fibre Formation

To achieve high-density cell-laden fibres, cells are incorporated into the initial collagen solution and subjected to interfacial spinning (Fig. 3a; Extended Data Fig. 4a,b).

**Fig. 3.**
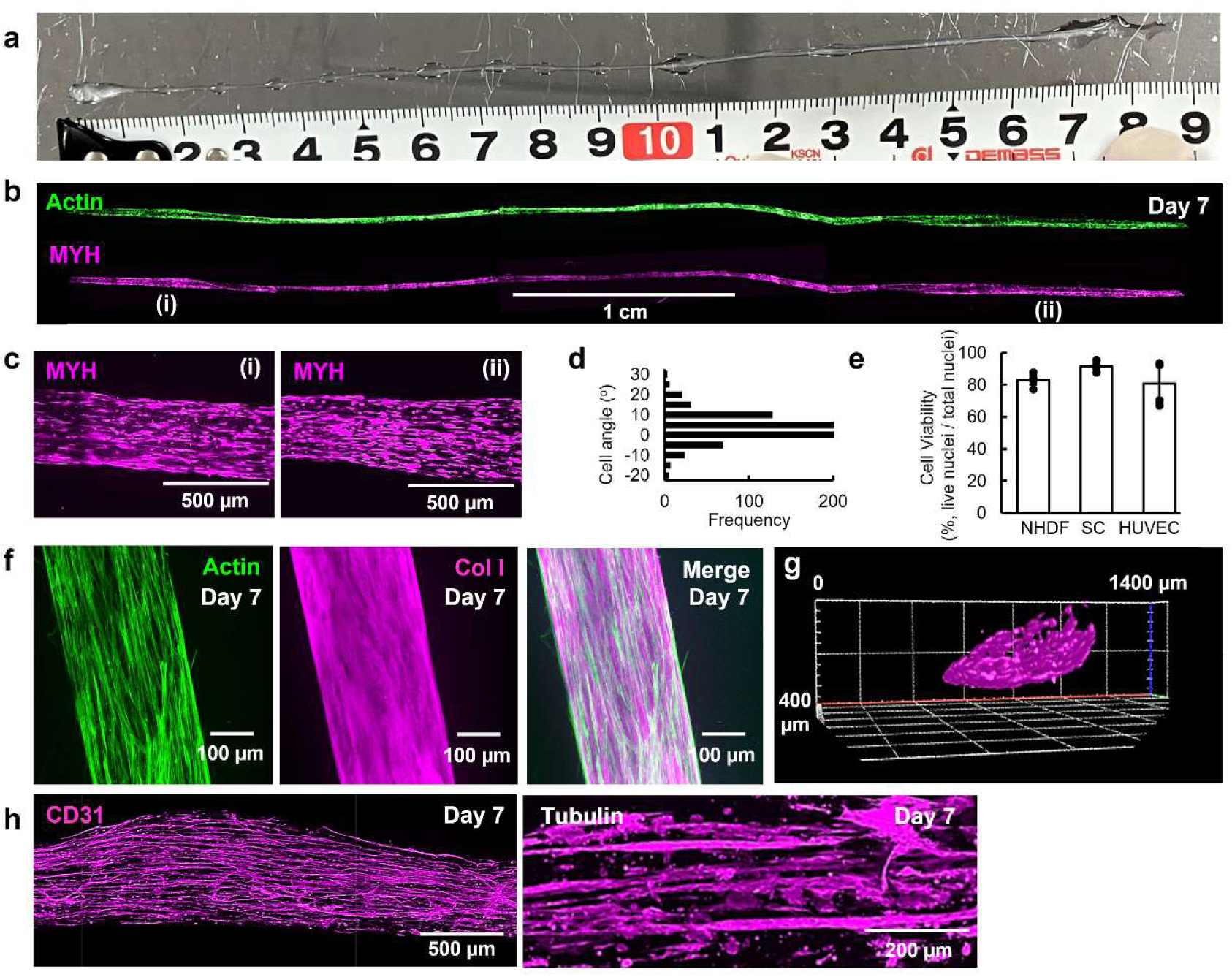
Fabrication and characterization of centimetre-scale cell-laden collagen fibres. a, Macroscopic photograph of SC-laden fibres alongside a ruler. **b,** Stitched immunofluorescence images of 5 cm SC fibres stained with phalloidin (green) and anti-myosin heavy chain (MYH) antibody (magenta). **c,** High-magnification images of regions (i) and (ii) indicated in **b**. **d,** Histogram of cell orientation angles calculated from MYH-immunostained SCs in Extended Data Fig. 4d (total fibres counted = 958). Angles represent deviation from the longitudinal axis (0°). **e,** Cell viability of NHDF-, SC-, and HUVEC-laden fibres (*n* = 3). Viability was calculated as the total number of DAPI-stained nuclei minus ethidium homodimer-positive (dead) cells. **f,** Immunofluorescence images of NHDF-laden fibres stained with phalloidin (green) and anti-collagen I antibody (magenta). **g,** 3D reconstructed images from z-stacked immunofluorescence of SC-laden fibres stained with anti-MYH antibody. **h,** Immunofluorescence images of HUVEC-and SH-SY5Y-laden fibres stained with anti-CD31 (HUVEC) and βIII-tubulin (SH-SY5Y) antibodies, respectively.

A notable feature is the spontaneous self-organization of encapsulated cells within the fibres (Supplementary Video 5). Cells adhere to the fibre interior, where they proliferate and align along the longitudinal axis (Extended Data Fig. 4c). Immunofluorescence imaging after culture reveals that satellite cells (SCs) are highly aligned along the fibre axis (0°) across the entire length, exhibiting elongated morphologies with well-developed F-actin filaments at higher magnification (Fig. 3b,c, Extended Data Fig. 4d). Quantitative analysis of orientation angles shows that most cells align within ±10° of the longitudinal axis (Fig. 3d).

This suggests that hierarchical collagen bundles within the fibre (Fig. 1) serve as robust contact guidance cues that direct cellular alignment at the macroscopic scale. Furthermore, this process induces minimal cytotoxicity in encapsulated cells. Assessments using normal human dermal fibroblasts (NHDFs), SCs and human umbilical vein endothelial cells (HUVECs) show high viability immediately after spinning (Fig. 3e; Extended Data Fig. 4e,f), with cellular activity and spreading maintained over 7 days. Across cell densities ranging from 1.0 × 10^6^ to 6.0 × 10^6^ cells ml^−1^ (Extended Data Fig. 5a,b), increasing density reduces fibre length (Extended Data Fig. 5c), while high viability (Extended Data Fig. 5d) and expression of functional proteins, including MYH4, are maintained. As continuous formation of macroscopic fibres is essential for scalable tissue engineering, an initial cell density of 1.0 × 10^6^ cells ml^−1^ is used as the standard condition. Even at this baseline density, encapsulated cells exhibit robust proliferation, with a marked increase in total cell number over the culture period (Extended Data Fig. 5e,f). Confocal microscopy of NHDF-laden fibres confirms a homogeneous cellular distribution throughout the construct, extending from the surface to the fibre core (Fig. 3f,g; Supplementary Video 6). Such uniform three-dimensional distribution is difficult to achieve using conventional post-seeding approaches, establishing this system as a uniaxially aligned tissue in which cells proliferate and form interconnected networks from within^20^. Consequently, this approach provides a versatile platform for engineering tissues in which structural anisotropy underpins functional expression, including muscle, vascular and neural tissues. Consistent with this, tissue-specific functional proteins, such as tubulin networks in neuronal cells (SH-SY5Y), CD31 in HUVECs (Fig. 3h), as well as type I collagen (Col I) in NHDFs and MYH4 in SCs are expressed within the fibres. To accommodate the specific requirements of these diverse tissues, culture conditions and physical anchoring strategies are tailored for each construct (Extended Data Fig. 6). Furthermore, encapsulation of adipose-derived stem cells (ADSCs) generates adipose-like fibres with intracellular lipid accumulation (Supplementary Fig. 2). These results indicate that the platform provides not only topographical alignment but also enables customizable biochemical reconstitution of tissue-specific microenvironments.

### Multi-Omics Integration Reveals Pathological Mechanotransduction

To evaluate the functionality of cell-laden fibres, we aimed to elucidate the molecular mechanisms by which cells sense their three-dimensional environment and establish their phenotypes. To this end, we performed integrated multi-omics analyses combining transcriptomics (RNA-seq) and proteomics. Principal component analysis (PCA) revealed distinct clustering of transcriptomic and proteomic profiles according to culture method (2D, Ran, and Fib), indicating that the global molecular landscape is fundamentally reshaped by the physical microenvironment (Supplementary Fig. 3). Initial global profiling revealed substantial and distinct alterations in the number of differentially expressed genes (DEGs) and proteins (DEPs) across culture formats (Supplementary Fig. 4). Comparison between aligned tissue (Fib) and random gel (Ran) showed that cytoskeletal and sarcomere-related proteins (e.g., ACTN1, MYH9) were significantly upregulated in Fib (Fig. 4a,b; Extended Data Fig. 7a). This confirms that structural alignment promotes cellular organization via mechanotransduction (Extended Data Fig. 7b,c)^21–23^. Comparison with conventional 2D culture (Fib vs. 2D) revealed an intriguing divergence: in Fib, genes and proteins associated with the mitochondrial respiratory chain and muscle contraction were downregulated relative to 2D cultures. Conversely, factors associated with the “amorphous matrix,” including collagens (e.g., COL6A1) and glycoproteins, as well as ECM interaction factors, were markedly upregulated (Fig. 4c,d)^24,25^.

**Fig. 4.**
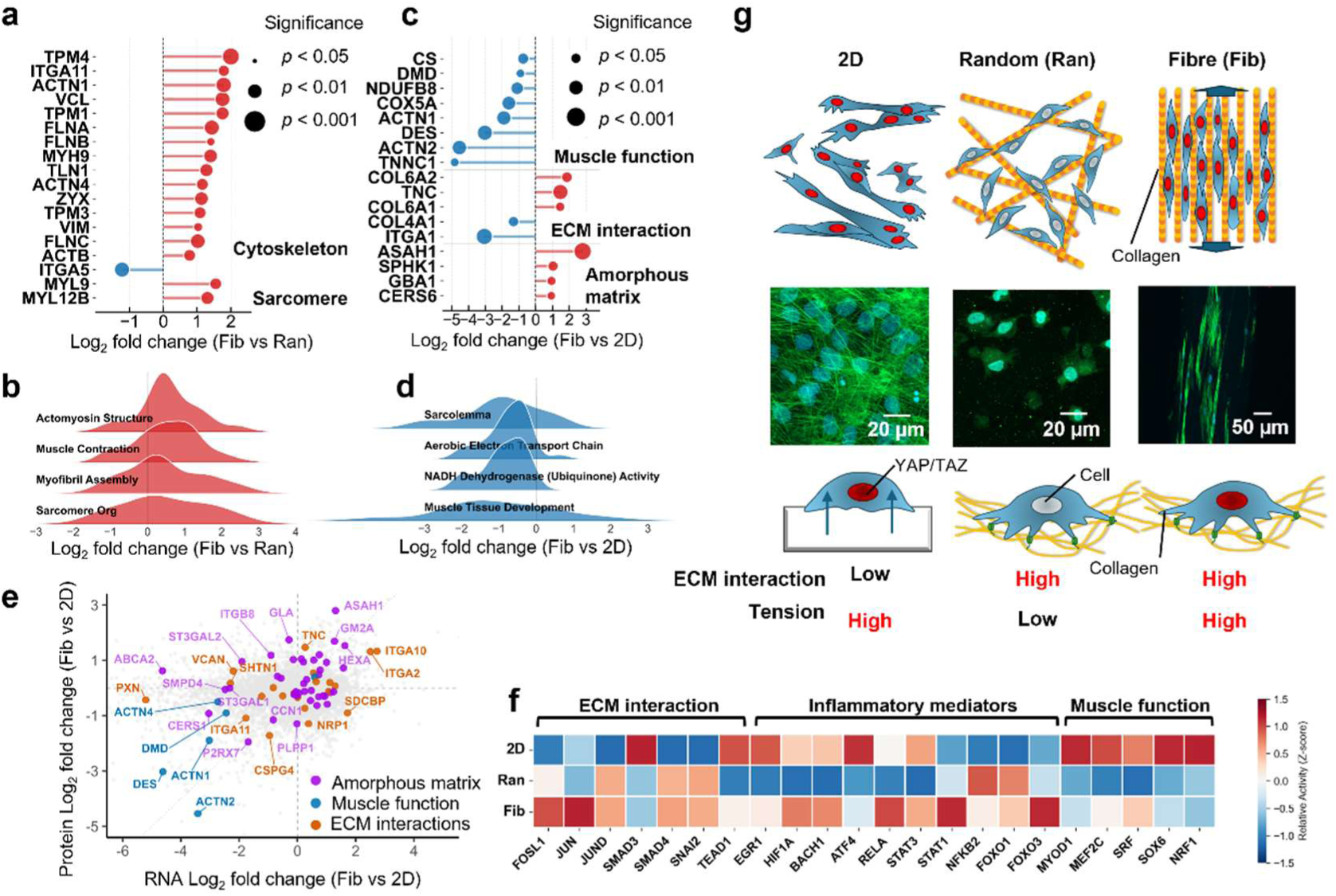
Multi-omics analysis of myoblast-laden fibres (Fib) compared with myoblasts in random collagen fibres (Ran) and on TCPS dishes (2D). a, Differentially expressed proteins (DEPs) related to the cytoskeleton and sarcomere in Fib compared with Ran. **b,** Ridge plot of gene set enrichment analysis (GSEA) for structural pathways in Fib versus Ran. **c,** DEPs related to muscle function, ECM interaction, and amorphous matrix in Fib compared with 2D. **d,** Ridge plot of GSEA for muscle-related pathways in Fib versus 2D. **e,** Transcriptome–proteome discordance scatter plot comparing log₂ fold changes of RNA and protein expression (Fib vs. 2D) (purple: amorphous matrix synthesis; blue: muscle function; orange: ECM interactions). **f,** Heatmap of inferred transcription factor (TF) activities based on Z-scores. **g,** Schematic illustration of proposed mechanotransduction mechanisms across 2D, Ran, and Fib environments, with confocal images of SCs stained with phalloidin and DAPI in each condition.

In the transcriptome–proteome correlation plot (Fig. 4e), muscle function–related factors (blue) clustered in the lower-left quadrant, reflecting pronounced downregulation of contractile elements in the Fib environment. This phenotypic shift is further supported by the accumulation of amorphous matrix–related factors (purple) in the upper-right quadrant, collectively indicating a coordinated transition from muscle maturation toward active ECM remodeling^26^. To explore the molecular basis of this shift, transcription factor (TF) activities were inferred, revealing three major clusters that fluctuated specifically under the Fib condition (Fig. 4f). Functionally, these clusters were classified into three groups: ECM interaction/fibrosis drivers^30,31^, inflammatory mediators^32–34^, and muscle function regulators^35,36^. Notably, the prominent activation of the first two groups (fibrosis and inflammation) indicates that the Fib environment induces not only muscle maturation but also a pathological state resembling fibrosis and inflammation in response to excessive mechanical stress^37^. Based on these findings, we propose a model of mechanotransduction in 3D engineered tissues (Fig. 4g). Unlike 2D culture (high tension, low ECM interaction) or Ran (low tension, high ECM interaction), the Fib environment uniquely exposes cells to both high ECM interaction and high cellular contractility. We hypothesize that this specific physical microenvironment synergistically activates AP-1 (FOSL1, JUN, JUND)^38^ and TGF-β (SMAD4, SNAI2)^39^ signaling pathways via mechanosensors such as YAP/TAZ^40,41^, thereby driving complex tissue remodeling that mirrors the pathological fibrotic progression characteristic of type 2 diabetic muscle, rather than promoting homeostatic maturation^42^. This finding demonstrates that our platform is valuable not only for constructing tissues for regenerative medicine but also for establishing unprecedented in vitro models of skeletal muscle fibrosis and myopathies induced by mechanical stress.

### System Versatility

The principal strength of this approach lies in its versatility: the ability to construct arbitrary macroscopic 3D architectures simply by modifying the interfacial topology. By slightly modifying the contact modality between acidic collagen and PAA, we generated not only “1D fibres” but also “2D sheets” via flat incubation, “0D capsules” via dropping, and hollow “1D tubes” via continuous injection—all using the same materials and underlying principles (Fig. 5a; Supplementary Video 7). Furthermore, integration with a microfluidic device enabled continuous generation of more complex topological architectures, such as necklace-like structures (Extended Data Fig. 8). Interfacial complexation proceeds continuously; for example, during sheet fabrication, macroscopic, time-dependent growth of the sheet thickness is clearly observed from 1 to 72 h (Supplementary Fig. 5). Furthermore, confocal fluorescence microscopy (maximum intensity projections and cross-sectional slices) confirmed that cells are encapsulated within the internal 3D spaces of tubes and capsules, achieving a homogeneous distribution comparable to that in the fibres (Fig. 5a). To elucidate the structural basis that enables these diverse macrostructures to maintain their integrity in aqueous environments, we evaluated the barrier function and permeability of the resulting interfacial collagen sheet (Fig. 5b). Strikingly, compared with conventional physically crosslinked bulk collagen gels, the interfacial film formed by our process exhibited markedly lower permeability coefficients (*P_e_*) for both low–molecular-weight (4 kDa) and high–molecular-weight (2,000 kDa) dextran molecules (Fig. 5b, middle; Supplementary Fig. 6a,b). Fluorescence microscopy further revealed a highly compact network structure (Fig. 5b, right), indicating that rapid, high-density electrostatic packing at the interface functions as a physical barrier. This barrier function was further supported by cellular behavior on the 2D sheets: owing to the dense network, cells did not infiltrate deeply into the matrix but instead predominantly organized along the tissue surface and available interstitial spaces (Fig. 5a). In conclusion, developed interfacial biofabrication technology embodies “ultimate simplicity”, which does not require complex equipment or chemical crosslinking and enables hierarchical structures comparable to native tissues, superior mechanical robustness, and excellent cytocompatibility. By enabling unrestricted combinations of desired geometries and cell types, this technology is applicable not only to muscle and blood vessels but also to encapsulated or tubular tissues, offering a powerful, universal platform for next-generation tissue engineering and fabrication of sophisticated in vitro pathological models.

**Fig. 5.**
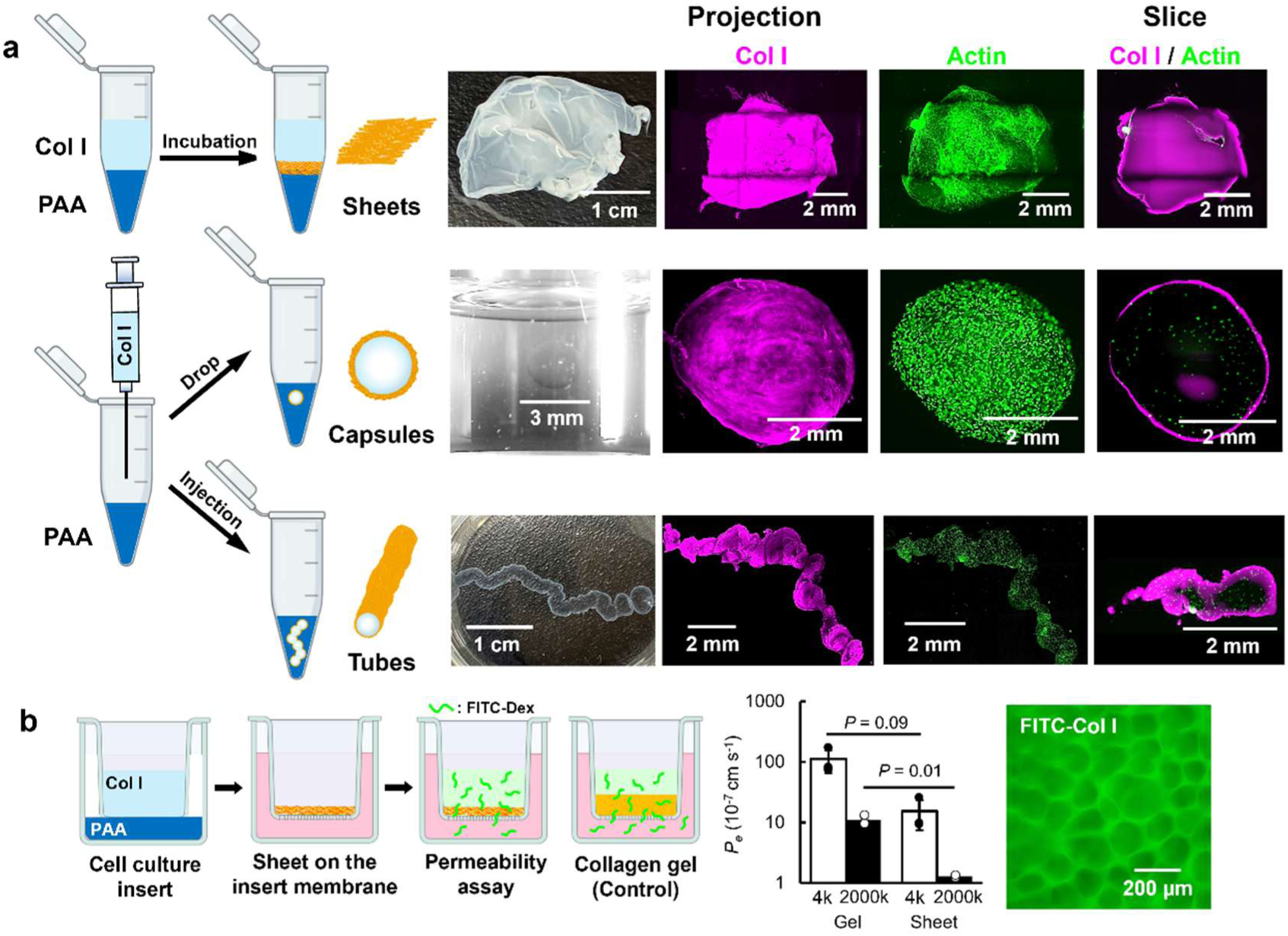
Fabrication of diverse milli-to centimetre-sized, dimension-controlled tissues and molecular permeability of 2D sheets. a, Schematic illustrations, macroscopic photographs, and light-sheet fluorescence images (maximum intensity projections and cross-sectional slices) of dimension-controlled tissues—2D sheets, 0D capsules, and 1D tubes—fabricated via interfacial collagen assembly (top), dropping (middle), and injection (bottom), respectively. Corresponding fluorescence images of cell-laden collagen constructs are stained with phalloidin (green) and anti-collagen I antibody (magenta). **b,** Permeability assay of the 2D sheet. Schematic illustration of the assay using a transwell insert (left). Permeability coefficients (*P_e_*) of conventional bulk collagen gel and the 2D sheet for 4 kDa FITC-dextran (Dex) and 2,000 kDa tetramethylrhodamine B isothiocyanate (TRITC)-Dex (middle) (*n* = 3). Fluorescence microscopy image of the FITC-labelled 2D sheet is shown on the right. Statistical significance was determined using an unpaired two-tailed Welch’s t-test.

## Conclusion

In summary, we established a biofabrication platform that leverages rapid interfacial polyion complexation between collagen and PAA to instantaneously spin highly aligned, pure collagen fibres. Overcoming longstanding trade-offs among mechanical robustness, cytocompatibility, and scalability, this method generates hierarchical architectures that mimic native ECM without the need for toxic chemical crosslinkers or complex equipment. Crucially, the uniaxially aligned topography serves as a robust contact guidance cue, enabling spontaneous self-organization and functional expression of diverse cell types, including muscle, vascular, and neural cells. Furthermore, integrated multi-omics analysis revealed that this unique 3D microenvironment—characterized by simultaneous high ECM interaction and cellular tension—activates distinct YAP/TAZ-mediated mechanotransduction pathways, offering unprecedented opportunities for modeling pathological states such as fibrosis. Beyond 1D fibres, the simplicity of this technology enables versatile construction of multidimensional architectures, including 2D sheets, 0D capsules, 1D tubes, and microfluidic-assisted necklace structures. This pristine, native-state collagen assembly platform bridges the gap between synthetic simplicity and biological complexity, offering substantial potential for advancing regenerative medicine, in vitro disease modeling, and scalable production of cultured meat.

## Materials and Methods

### Materials and Reagents

#### Polymers and Extracellular Matrices

Poly(acrylic acid) (PAA) with molecular weights (*M_W_*) of 5 kDa (165-18571, Fujifilm Wako), 25 kDa (165-18581, Fujifilm Wako), and 450 kDa (181285, Sigma-Aldrich) were used. Various types of collagen were evaluated to assess the effect of collagen source and organ origin. Type I, II, and III collagen are referred to as Col I, Col II, and Col III, respectively. The evaluated collagens included bovine skin-derived Col I (PSC-1-100-100, NIPPI), bovine cartilage-derived Col II (CL-22, Koken), bovine tendon-derived Col I (PSC-1-101-100, NIPPI), fish-derived Col I (PSC-1-500-100, NIPPI), and bovine skin-derived tropocollagen (Tropo, 50301, ibidi). Col III was purchased from NIPPI (PSC-3-100-20), and Col IV was purchased from Sigma-Aldrich (C5533). Laminin was also obtained from Sigma-Aldrich (L2020). For the polymer screening, the following materials were used: heparin sodium salt (H3393, Sigma-Aldrich), chondroitin sulfate A (C6737, Merck Millipore), chondroitin sulfate B (D3672, Tokyo Chemical Industry; TCI), chondroitin sulfate C (034-08801, Fujifilm Wako), chondroitin sulfate D (999999992301274, PG Research), chondroitin sulfate E (999999992301275, PG Research), hyaluronic acid (087-04511, Fujifilm Wako), sodium dextran sulfate (*M_r_* ≈ 40k, D8906, Merck Millipore), poly(styrenesulfonate) (*M_W_* = 70k, 041688.18, Thermo Fisher Scientific), fucoidan (20357, Funakoshi), λ-carrageenan (C3313, TCI), alginic acid sodium salt (Matsuba Chemical), poly(ethylene glycol) (95904, Sigma-Aldrich), poly(_L_-lysine) hydrochloride (*M_W_* = 15–30 kDa,P2658, Sigma-Aldrich) and poly(*γ*-glutamic acid) sodium salt (*M_W_* ≈ 400 kDa, 160-28592, Fujifilm Wako). Additional protein solutions included bovine serum albumin (BSA) (A3294, Sigma-Aldrich), human plasma fibronectin (F2006, Sigma-Aldrich) and gelatin (G9136, Sigma-Aldrich). BSA, fibronectin and gelatin were dissolved in phosphate-buffered saline (PBS) at a concentration of 5 mg mL^-1^ and subsequently diluted for the assay.

#### Cell Lines and Culture Reagents

Normal human dermal fibroblasts (NHDF, CC-2509, Lonza), human umbilical vein endothelial cells (HUVEC, CC2517A, Lonza), and the SH-SY5Y cell line (EC94030304-F0, KAC) were used. Bovine satellite cells (SC) and bovine adipose-derived stem cells (ADSC) were isolated according to previously reported methods^27^. Bovine tissues were obtained from a commercial abattoir (Sankyo Meat, Kagoshima, Japan) immediately after routine slaughter for commercial meat production. The donor animals were 27–29-month-old Japanese Black cattle, comprising one female (donor 71-5) and two steers (donors 72-2 and 73-4). Unless otherwise specified, all in vitro experiments using bovine satellite cells were performed using the cell line derived from donor 71-5. For omics analyses requiring biological triplicates, paired samples were prepared using the cell lines derived from all three independent donors (71-5, 72-2, and 73-4). For cell culture, Dulbecco’s Modified Eagle’s Medium (DMEM, 10569-010, Thermo Fisher Scientific), fibroblast growth factor (067-04031, Fujifilm), fetal bovine serum (FBS, 004-00025, Japan Bioserum), penicillin/streptomycin/amphotericin (P/S/A, 02892-54, Nacalai Tesque), Accumax (17087-54, Nacalai Tesque), and EGM™-2 MV BulletKit™ (CC3202, Lonza) were used for cell culture. Additional differentiation and culture supplements included a p38 inhibitor (SB203580, Selleck), retinoic acid (R2625-50MG, Sigma-Aldrich), nerve growth factor (141-07601, Fujifilm), oleic acid, and collagenase (038-22361, Fujifilm).

#### Chemicals, Assay Kits and Antibodies

For the permeability assay, 4 kDa FITC-dextran (FD4) and 2,000 kDa Tetramethyl rhodamine B-isothiocyanate (TRITC)-dextran (TD2000) were obtained from TdB Labs, and cell culture inserts (#3470) were purchased from Corning. For cell viability and proliferation assays, the LIVE/DEAD^®^ Viability/Cytotoxicity Kit (L3224, Thermo Fisher Scientific) and CellTiter-Glo^®^ 3D Cell Viability Assay (G9683, Promega) were used. Collagen quantification was performed using Direct Red 80 (365548, Sigma-Aldrich). Transcriptome analysis utilized the RNeasy Fibrous Tissue Mini Kit (#74704, Qiagen). For immunofluorescence staining, primary antibodies included anti-CD31 (MA3100, Thermo Fisher Scientific), anti-MYH4 (14-6503-82, Thermo Fisher Scientific), anti-COL1 (GTX26308, GeneTex) and anti-β-III tubulin (ab18207, Abcam). Secondary staining and counterstaining used normal goat serum (50062Z, Thermo Fisher Scientific), phalloidin (A12379, Thermo Fisher Scientific), DAPI (19178-91, Nacalai Tesque) and Hoechst 33258 (19173-41, Nacalai Tesque). Tissue clearing and mounting were performed using RapiClear 1.47 (RC147002, SUNJIN Lab) and RapiClear CS mounting gel (RCCS005, SUNJIN Lab). For fluorescence imaging of the collagen fibres, FITC-labeled type I collagen (K21, Collagen Research Center) and FITC-labeled PAA (PAA-FC-1, Nanocs Inc.) were used and mixed with the unlabeled collagen solution prior to spinning.

### Fabrication of Interfacial Collagen Fibres and 3D Structures

#### Basic Fibre Formation

To fabricate collagen fibres, a 100 µL aliquot of 30 mg mL⁻¹ PAA in PBS was added to a microcentrifuge tube. Then, 1,000 µL of 3 mg mL⁻¹ bovine skin-derived type I collagen solution was gently layered on top. Using fine tweezers, the PAA-collagen interface was scooped up and lifted slowly. The collected fibres were rinsed in ultrapure water.

#### Lower Limits for Fibre Formation

To determine the minimum concentrations required for interface formation, two setups were prepared: (1) 500 µL of a stepwise PBS-diluted series of 30 mg mL⁻¹ PAA (stained with blue food dye) was added to a tube and 500 µL of 3 mg mL⁻¹ collagen in acetic acid was carefully layered on top; (2) 500 µL of 30 mg mL⁻¹ PAA (stained blue) was added to a tube and 500 µL of stepwise diluted 3 mg mL⁻¹ collagen in 5 mM acetic acid was gently overlaid. Interface formation was continuously monitored for both conditions.

#### Optimization of Spinning Conditions

The conditions for fibre formation were optimized by altering the PAA solution parameters. *Effect of pH:* A 3 mg mL⁻¹ PAA (*M_W_* 25 kDa) solution in PBS was prepared and the pH was adjusted using 5 N NaOH or 1 N HCl. Fibres were spun against 3 mg mL⁻¹ bovine skin type I collagen and fibre lengths were measured with a ruler. *Effect of molecular weight:* 3 mg mL⁻¹ PAA solutions using 5 kDa, 25 kDa and 450 kDa polymers were tested against 3 mg mL⁻¹ bovine skin type I collagen. *Effect of concentration:* PAA (*M_W_* 25 kDa) solutions at 3 mg mL⁻¹ and 30 mg mL⁻¹ were tested against 3 mg mL⁻¹ bovine skin type I collagen. *Effect of collagen origin:* A 3 mg mL⁻¹ PAA (*M_W_* 25 kDa) solution was mixed with 3 mg mL⁻¹ of various collagens: bovine skin type Col I, III, bovine cartilage type Col II, fish Col I, bovine tendon Col I and bovine skin tropocollagen Col I.

#### Fabrication of Capsules, Tubes and Sheets

Three diverse macroscopic tissue architectures were produced as follows: *Capsules:* A PAA bath was prepared by adding 10 mL of 30 mg mL⁻¹ PAA in PBS to a 100 mm culture dish. 1.0 × 10^6^ ADSC were suspended in the collagen (3 mg mL⁻¹) solution and extruded intermittently into the PAA bath with a pipette, generating discontinuous droplets. After 60 s of gelation at room temperature, the spheres were transferred with a spatula to pre-warmed DMEM containing 10% FBS and cultured for 3 days under static conditions. *Tubes:* 100 µL of the cell-laden collagen solution was swiftly expelled into the air just above the PAA bath while the pipette was moved horizontally, generating a continuous strand on the bath surface. After 60 s of gelation, the PAA solution was carefully aspirated and replaced with pre-warmed DMEM containing 10% FBS. Constructs were incubated overnight. *Sheets:* 1 mL of a collagen solution (3 mg mL⁻¹ in 5 mM acetic acid) was dispensed into a vial. An equal volume of 30 mg mL⁻¹ PAA solution in PBS was carefully overlaid without disturbing the layer. The vial was kept undisturbed at 4 °C for 72 h. The resulting interfacial film was gently retrieved, transferred to a culture dish, covered with DMEM containing 10% FBS and seeded with 1.0 × 10^6^ NHDF.

### Microfluidic Device Fabrication and Generation of Topological Structures

To fabricate the microfluidic chip, photolithography was used to design a master mold on a silicon wafer containing a cross-junction with three inlets, one outlet, and 80-μm channels. Polydimethylsiloxane (PDMS) and its curing agent (Sylgard 184 Elastomer Kit) were mixed at a 10:1 weight ratio, poured into the master mold, and cured overnight at 60 °C. The PDMS replica was then peeled from the mold, and inlet and outlet ports were created using a biopsy punch (0.75 mm in diameter). The PDMS chip and a glass slide were sonicated in isopropanol for 5 min, sequentially washed three times with isopropanol and deionized water, and dried overnight at 60 °C. The surfaces were activated via oxygen plasma treatment to permanently bond the PDMS chip to the glass slide. To render the microchannels hydrophobic following the plasma treatment, the channels were coated with Aquapel®, which was flushed out after 2 min.

To continuously generate necklace-like topological structures, a polydimethylsiloxane (PDMS) microfluidic device with a cross-junction was employed. The dispersed phases consisted of the collagen solution and the PAA solution, while a continuous fluorinated oil phase (HFE 7500) was used to segment the flow and shear the droplets. The flow rates for the collagen solution, PAA solution and the continuous oil phase were precisely controlled using syringe pumps at 150, 150 and 500 µL h⁻¹, respectively.

### Structural and Morphological Characterization

#### Imaging and Morphology

The freshly collected fibres were immediately imaged under a phase-contrast microscope (IX71, EVIDENT) and macroscopic fibre lengths were measured with a ruler. High-speed imaging of the fabrication process was performed using a high-definition high-speed camera (HAS-EX, DITECT) at a frame rate of 1,000 fps. To observe the dried morphology, washed collagen fibres were vacuum-dried overnight at room temperature.

#### Scanning Electron Microscopy (SEM)

For SEM analysis, dried samples were sputter-coated with osmium using an osmium coater (Vacuum Device) and imaged at an accelerating voltage of 5.0 kV using an ultra-high-resolution scanning electron microscope (SU9000, Hitachi-High Technologies).

#### Atomic Force Microscopy (AFM)

Surface topography and microfibril structures of the dried fibres were characterized using an atomic force microscope (SPM-Nanoa, SHIMADZU) operated in contact mode. The deflection error signals were recorded to clearly visualize the microfibril edges for subsequent pitch measurement.

#### Image Analysis and Quantification

The orientation angles of the collagen bundles and encapsulated cells relative to the longitudinal axis of the macroscopic fibre were quantified using ImageJ software (NIH). For cell-free fibres, the fibrillar structures were extracted using the Ridge Detection plugin, whereas the orientation of the encapsulated cells from immunofluorescence images was analyzed by extracting their long axes using the Skeletonize function.

### Physicochemical and Mechanical Analysis

#### Turbidimetric Analysis of Complexation Kinetics

The kinetics of the collagen-PAA interfacial assembly were monitored turbidimetrically. Briefly, 100 µL of collagen solutions at varying concentrations (0–10 mg mL⁻¹) were added to a 96-well plate. A 30 mg mL⁻¹ PAA solution (*M_W_* 5 kDa) in PBS was then gently layered on top. The reaction kinetics were continuously tracked by measuring the absorbance (turbidity) at 600 nm every 1 min using a microplate reader at room temperature.

#### Crystal Structures and Chemical Composition Analysis

For X-ray diffraction (XRD), the spun fibres were washed with ultrapure water and vacuum-dried overnight at room temperature. Crystal structures were evaluated using an XRD instrument (AERIS, Malvern Panalytical). For Fourier Transform Infrared Spectroscopy (FT-IR) and Thermogravimetric Analysis (TGA), fibres were washed with DMEM containing 10% FBS overnight, followed by ultrapure water and then vacuum-dried overnight at room temperature. Chemical structures were analyzed via FT-IR (FT-720, HORIBA) in transmission mode (650–4,000 cm⁻¹, resolution = 2 cm⁻¹, 64 scans) at ambient temperature and pressure. TGA (STA7200RV, Hitachi High-Tech) was conducted under a nitrogen atmosphere flowing at 200 mL min⁻¹, with the sample placed in a platinum pan and heated at a constant rate of 10 °C min⁻¹.

#### Tensile Strength

Spun fibres were washed with ultrapure water and vacuum-dried overnight at room temperature. The fibre diameter was measured prior to testing. Samples were cut to a 30 mm length with scissors and the exact length was measured using calipers (Digimatic Caliper CD-15CP, Mitutoyo). Tensile tests (EZ-Test, SHIMADZU) were conducted at a speed of 1 mm min⁻¹. Fracture strength (MPa) and maximum stroke strain (%) at break were recorded. Young’s modulus (GPa) was calculated from the slope of the linear elastic region and toughness (kJ m⁻³) was determined from the area under the stress-strain curve at fracture.

#### Collagen Quantification Assay

To prepare samples for quantification, 100 µL of a 1 mg mL⁻¹ collagen solution diluted in acetic acid was added to each well of a 96-well plate. Subsequently, 100 µL of a 3 mg mL⁻¹ polymer solution was gently layered on top and the plate was incubated overnight. The supernatant was carefully aspirated, avoiding disturbance to the sedimented membrane. Wells were washed with ultrapure water and dried completely overnight at 37 °C. The dried samples were washed with distilled water and stained with 0.1% Direct Red 80 in distilled water containing 1% acetic acid. The bound dye was eluted with 100 µL of 0.1 M NaOH and absorbance was measured at 540 nm using a microplate reader (Infinite200PRO M Plex, Tecan).

#### Rheology

Fibres were spun using a 3 mg mL⁻¹ collagen solution and a 30 mg mL⁻¹ PAA (*M_W_* 5 kDa) solution in PBS, then washed with ultrapure water prior to measurement. As a control, a bulk collagen hydrogel was prepared by mixing the 3 mg mL⁻¹ collagen solution with a neutralizing buffer (10 × PBS and 0.05 M NaOH, 1:1 v/v) at a 4:1 ratio on ice. The mixture (300 µL) was quickly transferred into a cell culture insert and incubated overnight at 37 °C for complete gelation. Rheological properties were evaluated using a rheometer (MCR302, Anton Paar) equipped with a 25 mm parallel plate geometry at room temperature. Strain-dependent viscoelastic properties were measured at a fixed frequency (*f*) of 1 Hz.

#### Permeability Assay

To prepare the PAA-collagen interfacial film, 100 µL of a 3 mg mL⁻¹ collagen solution was added to a cell culture insert (Corning). The insert was placed into a 24-well plate containing 1 mL of a 30 mg mL⁻¹ PAA solution in PBS, allowing the two solutions to contact via the membrane and form an interfacial film overnight at 4 °C. For the control collagen hydrogel, 100 µL of the collagen solution was mixed with 25 µL of neutralizing buffer (25 mM HEPES and 0.05 M NaOH, 1:1 v/v) within the insert, while the reservoir was filled with 1 mL of PBS, followed by overnight incubation at 37 °C. An empty insert equilibrated with PBS was used as a blank control. Fluorescent tracers (4 kDa FITC-dextran and 2,000 kDa TRITC-dextran) were dissolved in PBS at 0.5 mg mL⁻¹ and applied to the apical side (inside the insert). At predetermined time points (1, 5, 15, 30, 45, 60, 75, 90, 105, 120 and 180 min), 50 µL aliquots were collected from the reservoir and immediately replenished with an equal volume of fresh PBS. Fluorescence intensity was measured using a microplate reader. To account for the dilution effect caused by buffer replenishment, the cumulative amount of permeated dextran (*Q_n_*, mg) at each time point *n* was corrected using Equation 1:

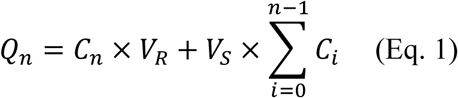

where 𝐶_n_ is the measured concentration in the reservoir (mg mL⁻¹), 𝑉_R_ is the total volume of the reservoir (mL) and 𝑉_S_ is the sampling volume (mL). The apparent permeability coefficient *P_app_* was calculated from the steady-state flux using Equation 2:

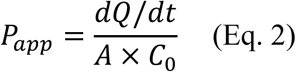

where 𝑑𝑄/𝑑𝑡 is the permeation rate (mg s⁻¹) determined from the linear slope of the *Q_n_* versus time plot, 𝐴 is the effective surface area of the membrane (cm²) and 𝐶_O_ is the initial concentration of dextran in the donor chamber (mg mL⁻¹). Finally, the effective permeability coefficient (𝑃_e_) of the collagen-PAA film or collagen gel was determined by subtracting the resistance of the blank control membrane using Equation 3:

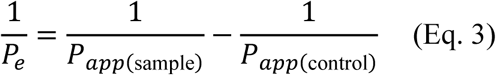

### Cell Culture and 3D Encapsulation

#### Cell Culture

NHDF, SH-SY5Y and ADSC were pre-cultured in DMEM containing 10% FBS and 1% P/S/A. HUVEC were cultured using EGM-2 MV BulletKit medium in a 5% CO₂ incubator at 37 °C. SC were cultured for proliferation in BM#2, consisting of high-glucose DMEM containing 1% P/S/A, 20% FBS, 4 ng mL⁻¹ fibroblast growth factor and 10 µM p38 inhibitor. To induce differentiation, ADSC were cultured in DMEM supplemented with 10% FBS and 100 mM oleic acid, while SC were differentiated in DMEM with 2% FBS. For SH-SY5Y cells, differentiation was initiated in DMEM containing 2% FBS and 10 µM retinoic acid, followed by maturation in DMEM supplemented with 1% FBS and 10 µM nerve growth factor. All cells were pre-cultured and differentiated at 37 °C in a 5% CO₂ incubator.

#### Cell Fibre Fabrication

SC (1.0 × 10^6^ cells) were suspended in 1,000 µL of 3 mg mL⁻¹ bovine skin collagen on ice. Separately, PAA (*M_W_* 5 kDa) was dissolved to 30 mg mL⁻¹ in PBS and aliquoted into a sterile 1.5 mL microtube. The collagen-cell suspension was gently layered atop the PAA solution. Immediately after layering, the interface was captured with sterile fine-tip forceps and slowly lifted to generate a continuous cell-laden fibre. For the NHDF, HUVEC, co-culture (NHDF + HUVEC) vessel and SH-SY5Y nerve fibres, the total cell concentration was kept identical to that used for the SC fibres. Following fabrication, each fibre was transferred to a 6-well plate containing 3 mL of pre-warmed proliferation medium and incubated at 37 °C with 5% CO₂. For the vascular fibres, both ends of the construct were anchored to the well walls using fibrin gel to prevent macroscopic shrinkage during culture. The subsequent culture schedules were tailored for each specific cell type to optimize tissue formation:

NHDF fibres: Cultured in proliferation medium (DMEM supplemented with 10% FBS) for 5 days.

SC fibres: Cultured in proliferation medium for 5 days, followed by 14 days in differentiation medium.

Nerve fibres (SH-SY5Y)

Maintained in proliferation medium for 1 day, shifted to differentiation medium for 1 day and subsequently matured for 6 days.

Vessel fibres (NHDF + HUVEC)

Cultured in proliferation medium for 7 days. At the end of their respective culture periods, all fibres were fixed in 4% paraformaldehyde for subsequent analyses. To evaluate the impact of initial cell density, the procedure was repeated using 1.0 × 10^6^, 3.0 × 10^6^, or 6.0 × 10^6^ SC.

### Viability and Immunofluorescence Staining

#### Cell Number and Viability Analysis

For viability analysis, a staining solution was prepared by adding Calcein AM, Ethidium Homodimer-1 (EthD-1) and 5 µL Hoechst 33258 to phenol-red-free medium supplemented with 10% FBS. Samples were incubated at 37 °C for 3 h, washed with PBS, mounted on glass slides with a drop of PBS and covered with a coverslip. Fluorescence images were acquired (IX71, EVIDENT) and viability was calculated as follows:

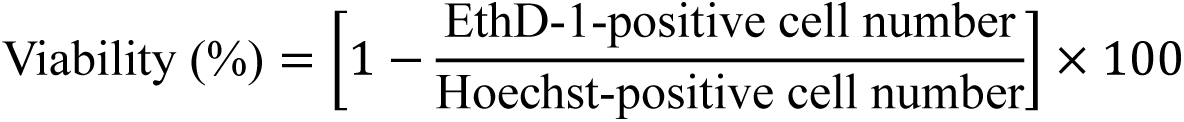

To quantify cell numbers, fibres were cut into 10 mm segments and one segment was placed per well of a 96-well plate containing 100 µL of culture medium. ATP content was measured using the CellTiter-Glo 3D Cell Viability Assay and absolute cell numbers were calculated from a standard curve.

#### Immunostaining and Optical Clearing

Fibres were carefully extended and fixed in 4% paraformaldehyde for 3 h at room temperature. Samples were permeabilized and blocked in PBS containing 1% normal goat serum and 0.2% Triton X-100 for 1 h at room temperature. Primary antibodies were diluted in the blocking buffer and incubated with the fibres overnight at 4 °C. After washing with PBS, specimens were incubated overnight at 4 °C with fluorophore-conjugated secondary antibodies, phalloidin and DAPI. Stained samples were immersed in 1 mL of RapiClear 1.47, protected from light and gently inverted overnight at room temperature. Finally, RapiClear CS mounting gel was heated to 75 °C, poured into a mold around the sample and allowed to solidify. Fluorescence images were acquired using a confocal laser scanning microscope (LSM910, Zeiss) and a light-sheet fluorescence microscope (Lightsheet-7, Zeiss). Images were processed and analyzed using Zen software and ImageJ.

### Transcriptomics and Proteomics Analysis

#### Transcriptome Analysis

SC-loaded fibres (Fib) were fabricated using collagen and PAA. Fibres were cultured in BM#2 for 5 days (proliferation) and then switched to differentiation medium for 14 days. Random 3D constructs (Ran) were generated by gently pipetting the cell-laden collagen suspension with the PAA solution and the resulting aggregates were cultured identically to the fibres. For 2D controls, SC were seeded in 6-well plates, grown for 2 days in proliferation medium and then for 14 days in differentiation medium. Total RNA was extracted using the RNeasy Fibrous Tissue Mini Kit. To ensure rigorous paired comparisons across all culture conditions (2D, Ran, and Fib) and omics platforms (transcriptomics and proteomics), the biological triplicates were strictly derived from the exact same three independent primary cell lines (strains 71-5, 72-2, and 73-4) established in this study.

#### Proteomic Analysis

Tissue samples (Fib, Ran, 2D) were washed, minced into 1 mm pieces and dissociated by incubation with 1 mL of Accumax at 37 °C for 5 min, followed by centrifugation (10,000 rpm, 3 min). The pellet was resuspended in 1 mL of 2 mg mL⁻¹ collagenase, incubated at 37 °C for 10 min with shaking and centrifuged again. The cell pellet was stored at −150 °C. Protein extraction, trypsin digestion and liquid chromatography-tandem mass spectrometry (LC-MS/MS) were performed by Kazusa Genome Technologies. Proteomic data processing was performed using DIA-NN software (v1.81).

#### Bioinformatics Analysis

Principal Component Analysis (PCA) for RNA-seq data was performed using Python. The raw count matrix was processed using *pandas*, normalized to Counts Per Million (CPM) and log2(CPM + 1) transformed. Data were standardized (mean = 0, variance = 1) using *scikit-learn* and analyzed to visualize sample distribution^28^. Differential expression analysis was performed using R (v4.3.0) via the *DESeq2* package^29^ with the design model ∼ Cellline + Method. Differentially Expressed Genes (DEGs) and Proteins (DEPs) were defined as having an adjusted *P* value (FDR) < 0.05 and an absolute log₂ fold change ≥ log_2_(1.5) (≈ 0.585). Gene Set Enrichment Analysis (GSEA) was evaluated using the *fgsea* package^30^ against GO and KEGG databases (FDR < 0.1). Upstream transcription factor (TF) activities were inferred from bulk RNA-seq data using the *decoupleR* package and the DoRothEA regulon network and normalized to Z-scores^31^. For transcriptome-proteome integration, log₂FC values from both datasets were plotted on a 2D scatter plot using the *ggplot2* and *ggrepel* packages^32^.

## Statistical Analysis

The specific statistical test used for each experiment is specified in the corresponding figure legend. A *P* value < 0.05 was considered statistically significant. In all cases, quantitative data were obtained from at least three independent experiments. Statistical analyses and visualizations were performed using Microsoft Excel and Python (v3.11.8) with *pandas*, *statsmodels* and *scikit-posthocs* packages. Dispersion and precision measures (mean and standard deviation) are specified in the figure legends.

## Acknowledgements

This work was supported by the Japan Society for the Promotion of Science (JSPS) KAKENHI Grant Numbers JP22H05141, JP25H01220, and JP22K21348; the JSPS Bilateral Joint Research Project Grant Number JPJSBP120252301; the Japan Science and Technology Agency (JST) Grant Number JPMJPF2009; and the New Energy and Industrial Technology Development Organization (NEDO) Grant Numbers 21W2K022, 23201377-0, and 23201347-0; and the Deutsche Forschungsgemeinschaft (DFG) in the framework of Priority Programme (SPP 2451) “Engineered Living Materials with Adaptive Functions” within the project 541298489. We acknowledge the NGS core facility at the Research Institute for Microbial Diseases of Osaka University for the sequencing and data analysis.

## Author Contributions

A.Y. and M.M. conceived the project and designed the experiments. A.Y. performed the majority of the experiments, analyzed the data, and wrote the original manuscript. K.H. performed the experiments regarding the nerve and vascular fibres. A.W. performed the experiments regarding the collagen capsules. Y.S. conducted the experiments related to the microfluidic device under the supervision of A.P. S.K. and M.M. supervised the overall project, administrated the research, and acquired funding. M.M., A.P., and S.K. reviewed and edited the manuscript. All authors discussed the results and approved the final manuscript.

## Data Availability/Code Availability

### Data Availability

The RNA-sequencing data generated in this study have been deposited in the DDBJ Sequence Read Archive (DRA) under the BioProject accession number PRJDB18471. The proteomics data have been deposited in the jPOST/ProteomeXchange Consortium via the jPOST Partner Repository under the accession numbers JPST004530 (jPOST) and PXD076302 (ProteomeXchange). All other data supporting the findings of this study are available within the paper and its Supplementary Information. Source data are provided with this paper. Any remaining raw data will be available from the corresponding author upon reasonable request.

### Code Availability

No previously unreported custom computer code or algorithm was used to generate the results in this study. All multi-omics data analyses were performed using standard functions of the widely used software packages and tools described in the Methods section. Custom data processing scripts are available from the corresponding author upon reasonable request.

---

**Extended Data Fig. 1.**
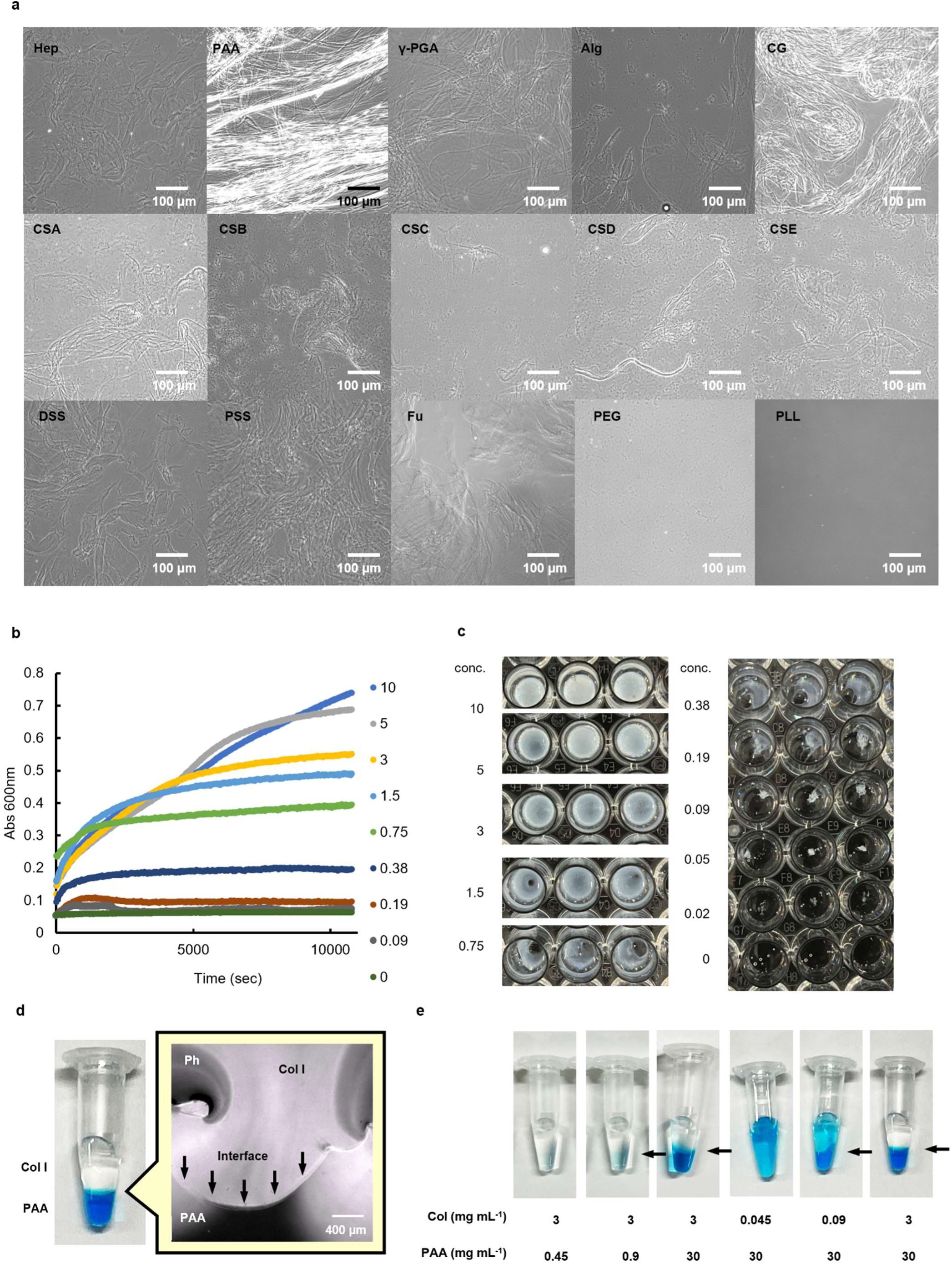
Physicochemical characterization and kinetics of collagen-polymer interfacial assembly. a, Phase-contrast microscopy images of collagen complexes formed with various polymers. Col I (3 mg mL^-1^) was mixed with various polymer solutions (3 mg mL^-1^ in PBS). Abbreviations: Hep, heparin; PAA, poly(acrylic acid); γ-PGA, poly(*γ*-glutamic acid); Alg, alginic acid; CG, carrageenan; CSA–CSC, chondroitin sulfate A–C; DSS, dextran sulfate; PSS, poly(styrenesulfonate); Fu, fucoidan; PEG, poly(ethylene glycol); PLL, poly(_L_-lysine). **b–c,** Kinetics of interface formation monitored by turbidimetric analysis. Time-course measurements of turbidity (absorbance at 600 nm) over 3 h (10,800 s) upon mixing varying concentrations of collagen (0 to 10 mg mL^-1^) (b) and the corresponding macroscopic appearance of the reaction mixtures in a multi-well plate (c). **d,** Photograph (left) and phase-contrast microscopy image (right) of the liquid-liquid interface formed between collagen (clear, top layer) and PAA solution (blue, bottom layer). Arrows indicate the solid-like interfacial membrane at the boundary. **e,** Macroscopic photographs of interface formation across varying concentrations of collagen and PAA. Collagen solutions and PAA solutions (stained with blue food coloring) were layered to determine the critical concentrations for stable film formation. Collagen and PAA solutions were prepared in 5 mM acetic acid and PBS, respectively. Arrows indicate successful interface formation.

**Extended Data Fig. 2.**
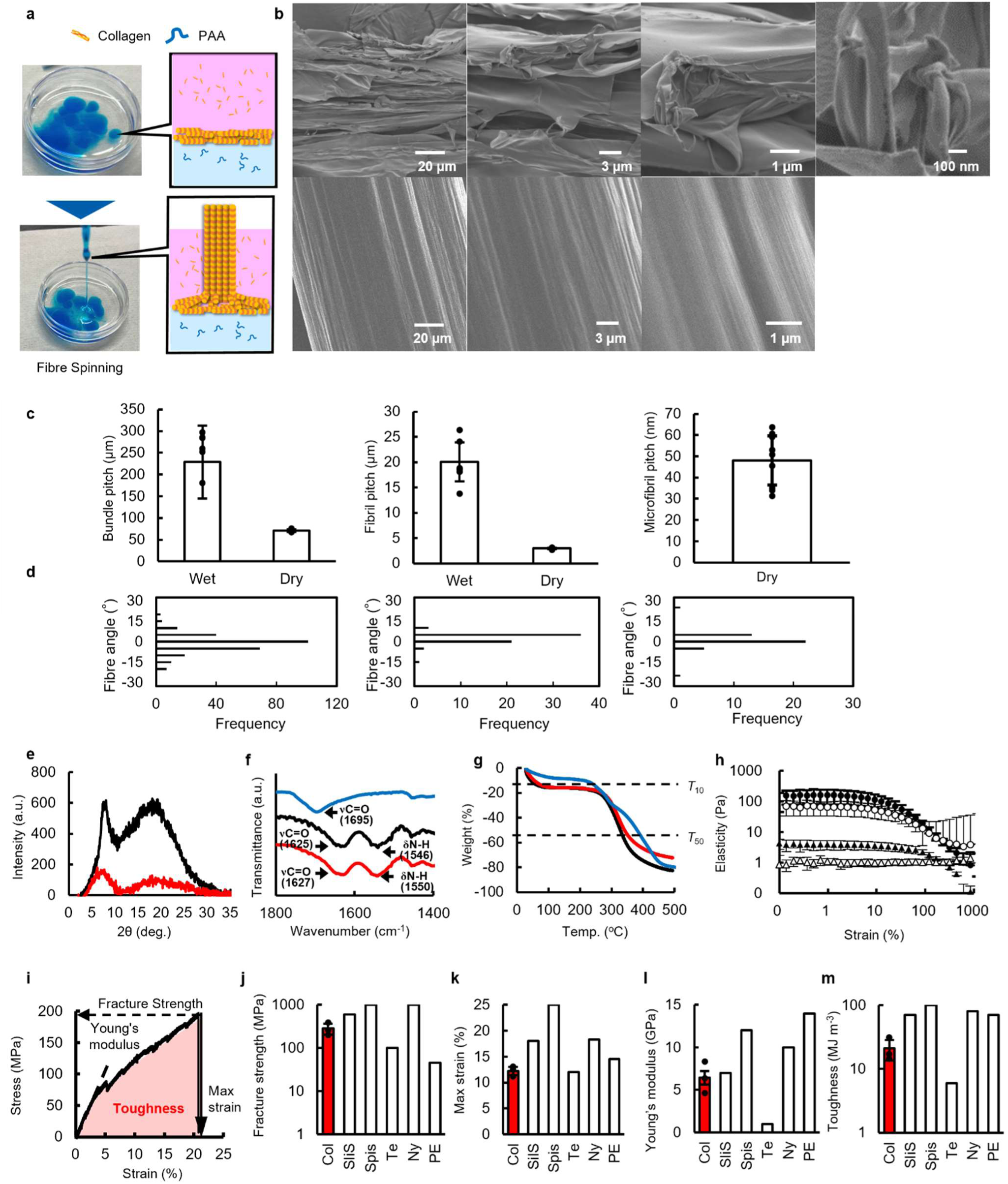
Structural and biomechanical characterization of uniaxially aligned collagen fibres. a, Photographs and schematic illustrations of the continuous fibre spinning process. **b,** Ultrastructural comparison between native collagen and fabricated fibres. Scanning electron microscopy (SEM) images at various magnifications of native collagen (lyophilized control, top) and fabricated PAA–collagen fibres (vacuum dried, bottom). The high-magnification SEM image corresponding to the rightmost panel is presented in Fig. 1b. **c,** Pitch evaluation of collagen bundles, fibrils, and microfibrils in wet and dry states. Error bars represent ± s.d. (*n* = 3–7 individual fibres per representative image). **d,** Histograms of fibre orientation angles measured from phase-contrast (left), atomic force microscopy (center), and SEM (right) images shown in Fig. 1b. Angles represent deviation from the longitudinal axis (0°). **e,** X-ray diffraction (XRD) patterns of native collagen (black) and fabricated fibre (red). **f,** Fourier transform infrared (FT–IR) spectra of native collagen (black), fibre (red), and PAA (blue). **g,** Thermogravimetric analysis (TGA) curves showing thermal degradation profiles of native collagen (black), fibre (red), and PAA (blue). *T*₁₀ and *T*₅₀ indicate temperatures at 10% and 50% weight loss, respectively. **h,** Dynamic viscoelastic properties (strain sweep) of the collagen film and gel. Storage (*G’*: closed symbols) and loss (*G’’*: open symbols) moduli are plotted against strain. Circles and triangles represent the film and gel, respectively (*f* = 1 Hz). **i,** Representative stress–strain curve of a dried interfacial collagen fibre, illustrating definitions of extracted mechanical parameters. The fibre was fabricated using bovine skin-derived type I collagen (3 mg mL⁻¹ in 5 mM acetic acid). **j–m,** Comparison of mechanical properties: fracture strength (j), maximum strain (k), Young’s modulus (l), and toughness (m) between fabricated collagen fibres (Col, red bars) and various natural and synthetic fibres (SilS, silkworm silk; SpiS, spider silk; Te, tendon collagen; Ny, nylon 6-6; PE, polyethylene). Error bars represent ± s.d. (*n* = 3 independent samples). Reference values were obtained from the literature. Tensile tests were performed at a crosshead speed of 1 mm min⁻¹.

**Extended Data Fig. 3.**
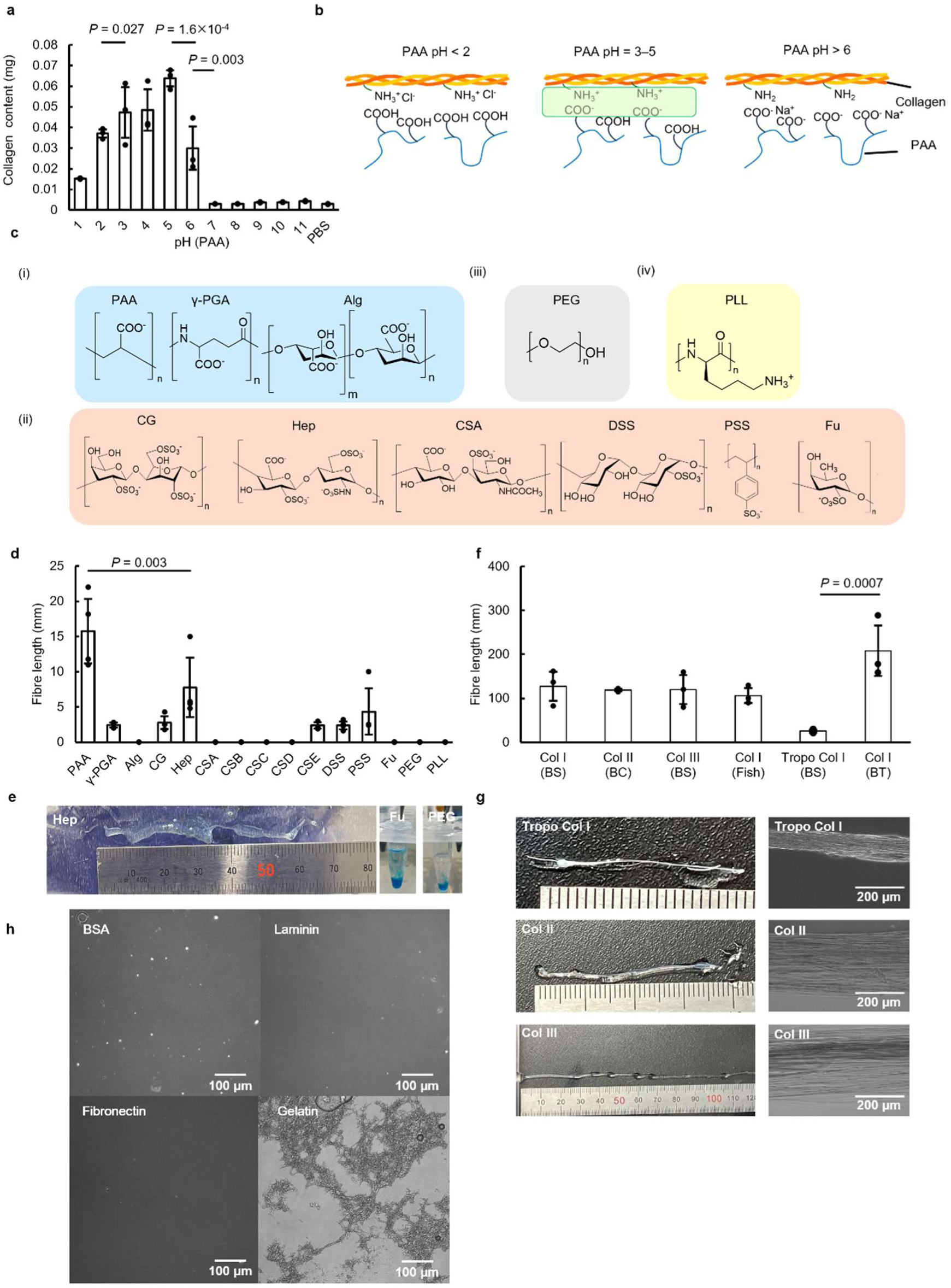
Electrostatic mechanisms and material scope of interfacial complexation. a, Quantitative analysis of collagen accumulation at the interface formed with PAA across different pH levels (pH 1–11), measured by Sirius Red staining. Error bars represent ± s.d. (*n* = 3 independent samples). Statistical significance was determined by one-way ANOVA with Tukey’s post hoc test. Exact *P* values are indicated in the graph. **b,** Schematic illustration of the electrostatic interaction model between collagen and PAA at different pH levels. **c,** Chemical structures of the polymer library classified by charge and functional groups: (i) carboxylated polyanions, (ii) sulfated or sulfonated polyanions, (iii) neutral polymer, and (iv) polycation. **d,** Effect of polymer type on fibre length. Abbreviations: PAA, poly(acrylic acid); γ-PGA, poly(*γ*-glutamic acid); Alg, alginic acid, CG, carrageenan, Hep, heparin; CSA–CSE, chondroitin sulfate A–E; DSS, dextran sulfate; PSS, poly(styrenesulfonate); Fu, fucoidan; PEG, poly(ethylene glycol); PLL, poly(_L_-lysine). Error bars represent ± s.d. (*n* = 3). Statistical significance was determined by one-way ANOVA followed by Tukey’s post hoc test. Exact *P* values are indicated in the graph. **e,** Macroscopic photographs of successful (Hep) and failed (Fu, PEG) fibre formations (polymer solutions were dyed with blue food coloring). **f**, Effect of collagen source and organ origin on fibre length. Col I, Col II, and Col III represent type I, II, and III collagen, respectively. BS: bovine skin, BC: bovine cartilage, BT: bovine tendon, Tropo: tropocollagen. Error bars represent ± s.d. (*n* = 3). Statistical significance was determined by one-way ANOVA followed by Tukey’s post hoc test. Exact *P* values are indicated in the graph. **g,** Macroscopic photographs and microscopy images of fibres composed of different collagen types. **h,** Phase-contrast microscopy images of mixtures containing PAA (molecular weight= 25 k Da, 3 mg mL^-1^) and various protein solutions (3 mg mL^-1^ BSA, laminin, fibronectin, and gelatin).

**Extended Data Fig. 4.**
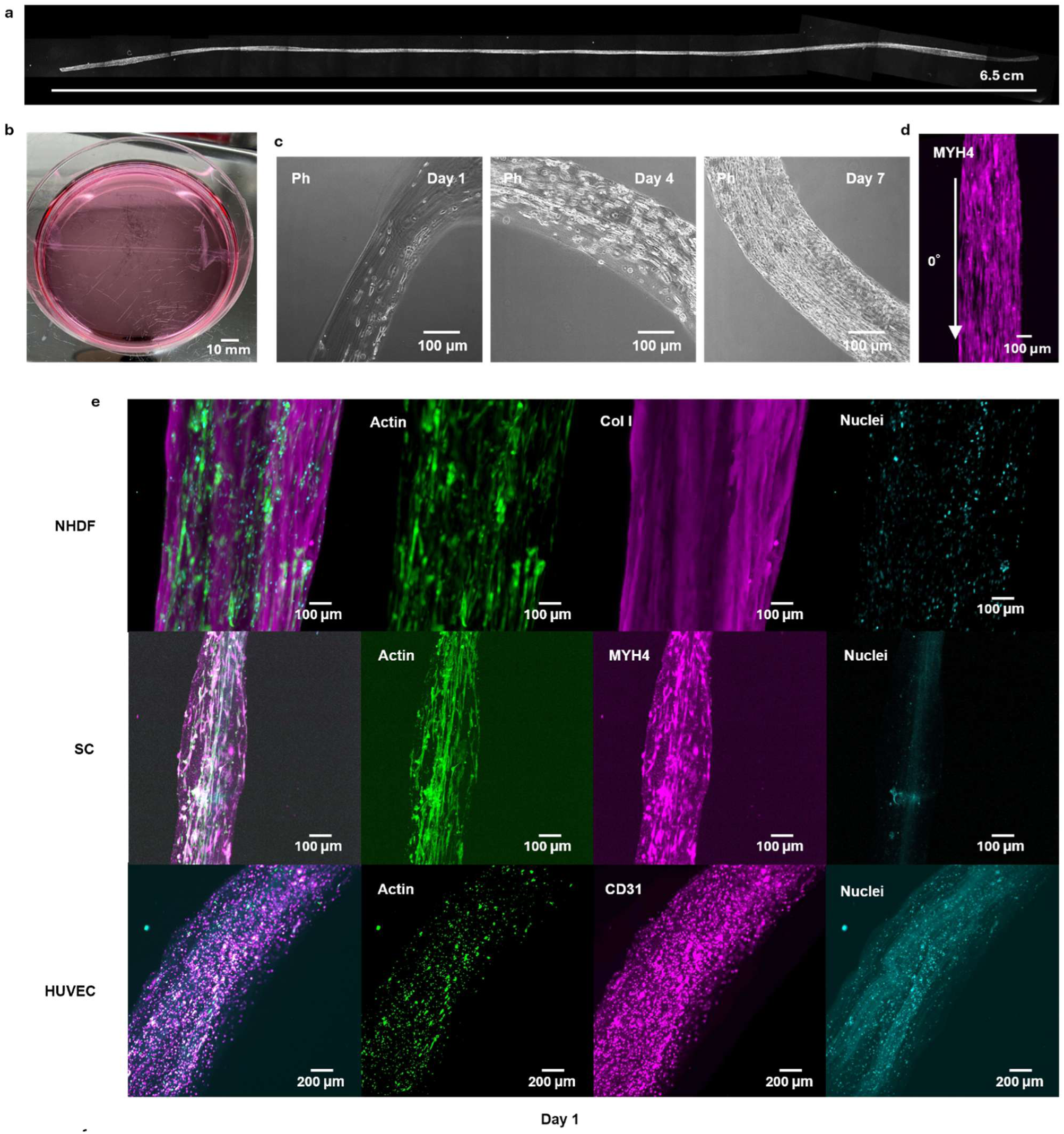
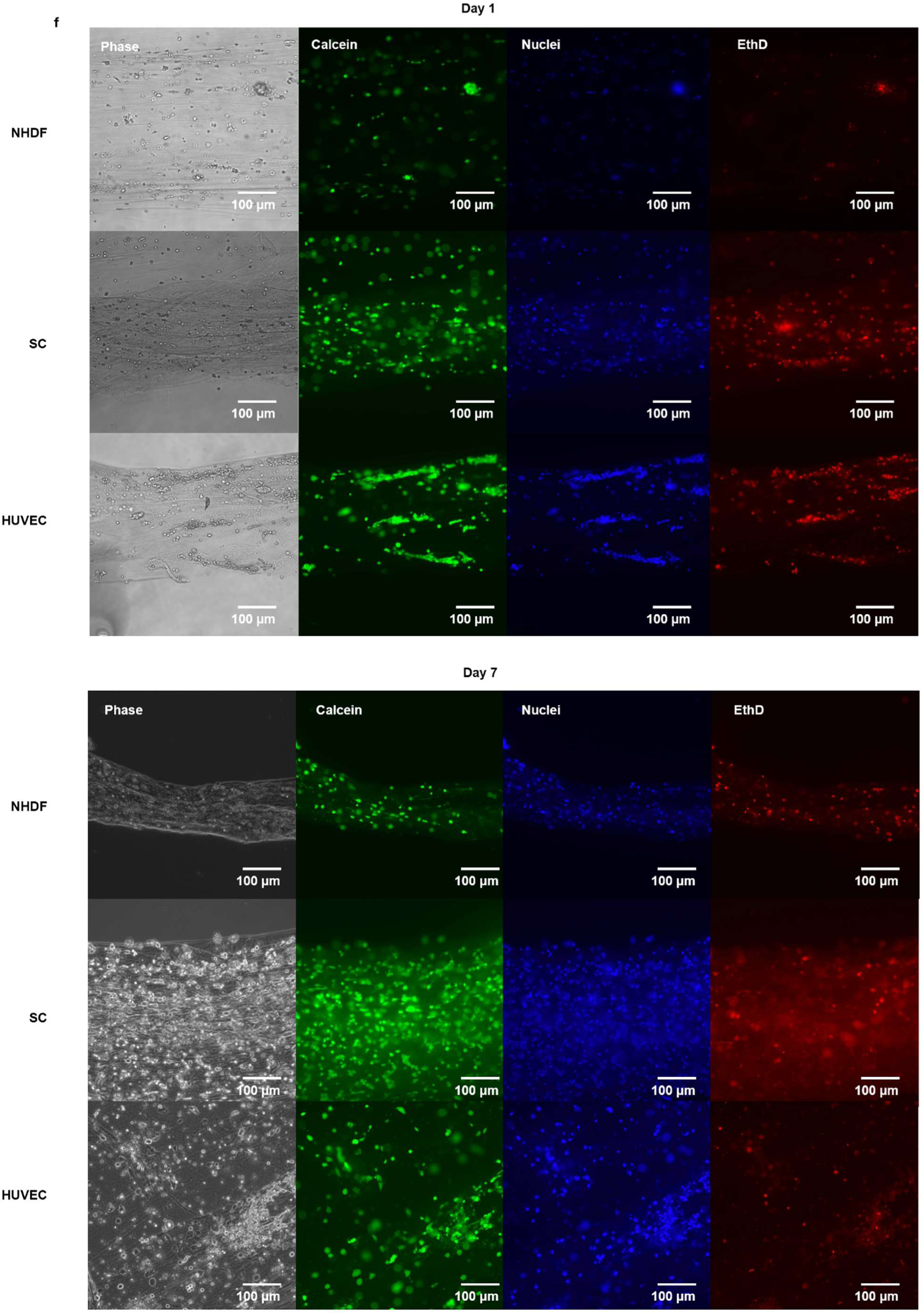
Cellular alignment and phenotypic expression within abricated fibres. a, Stitched phase-contrast image of an entire fibre. **b,** Macroscopic photograph of fibres floating in culture medium. **c,** Phase-contrast images showing time-dependent growth and alignment of SCs in the fibre at days 1, 4, and 7. **d,** Immunofluorescence image of the fibre stained for MYH4 (magenta), used for orientation analysis of filaments relative to the longitudinal axis (white arrow, 0°). **e,** Immunofluorescence images of fibres fabricated with NHDF, SC, and HUVEC. Cells were stained for specific markers (green: F-actin; blue: nuclei; magenta: Col I for NHDF, MYH4 for SC, and CD31 for HUVEC). NHDF and HUVEC fibres were cultured for 7 days; SC fibres were cultured for 7 days for proliferation and 5 days for differentiation. **f,** LIVE/DEAD fluorescence images of fibres prepared with different cell types (SC, HUVEC, and NHDF) from day 1 to day 7 (green: calcein AM; red: ethidium homodimer; blue: nuclei).

**Extended Data Fig. 5.**
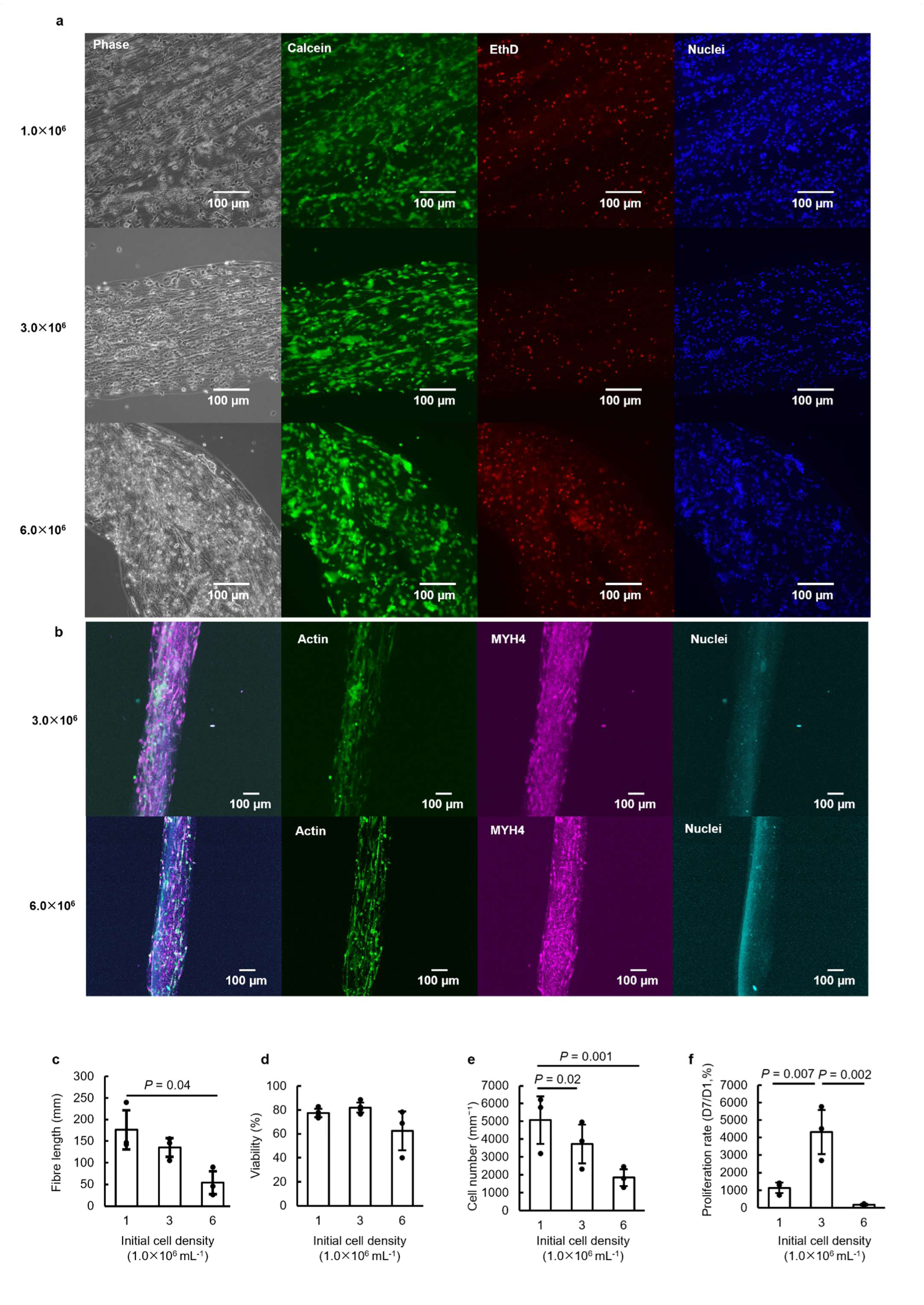
Optimization of initial cell density and long-term viability in 3D cell fibres. a, Phase-contrast and live–dead staining images of fibres fabricated at initial cell densities of 1.0, 3.0, and 6.0 × 10⁶ cells mL⁻¹ (green: calcein AM; red: ethidium homodimer; blue: nuclei). **b,** Immunofluorescence images of fibres fabricated at initial cell densities of 3.0 and 6.0 × 10⁶ cells mL⁻¹ (green: actin; magenta: MYH4; cyan: nuclei). **c–f,** Quantitative evaluation of fibres fabricated with varying initial cell densities (1, 3, and 6 × 10⁶ cells mL⁻¹): fibre length (c), cell viability at day 7 (d), cell number (e), and proliferation rate (f). Cell viability was calculated as the total number of DAPI-stained nuclei minus ethidium homodimer-positive (dead) cells. Proliferation rate was defined as the ratio of live cell number on day 7 to that on day 1. Error bars represent ± s.d. (*n* = 3 independent samples). Statistical significance was determined by one-way ANOVA with Tukey’s post hoc test. Exact P values are indicated in the graph.

**Extended Data Fig. 6.**
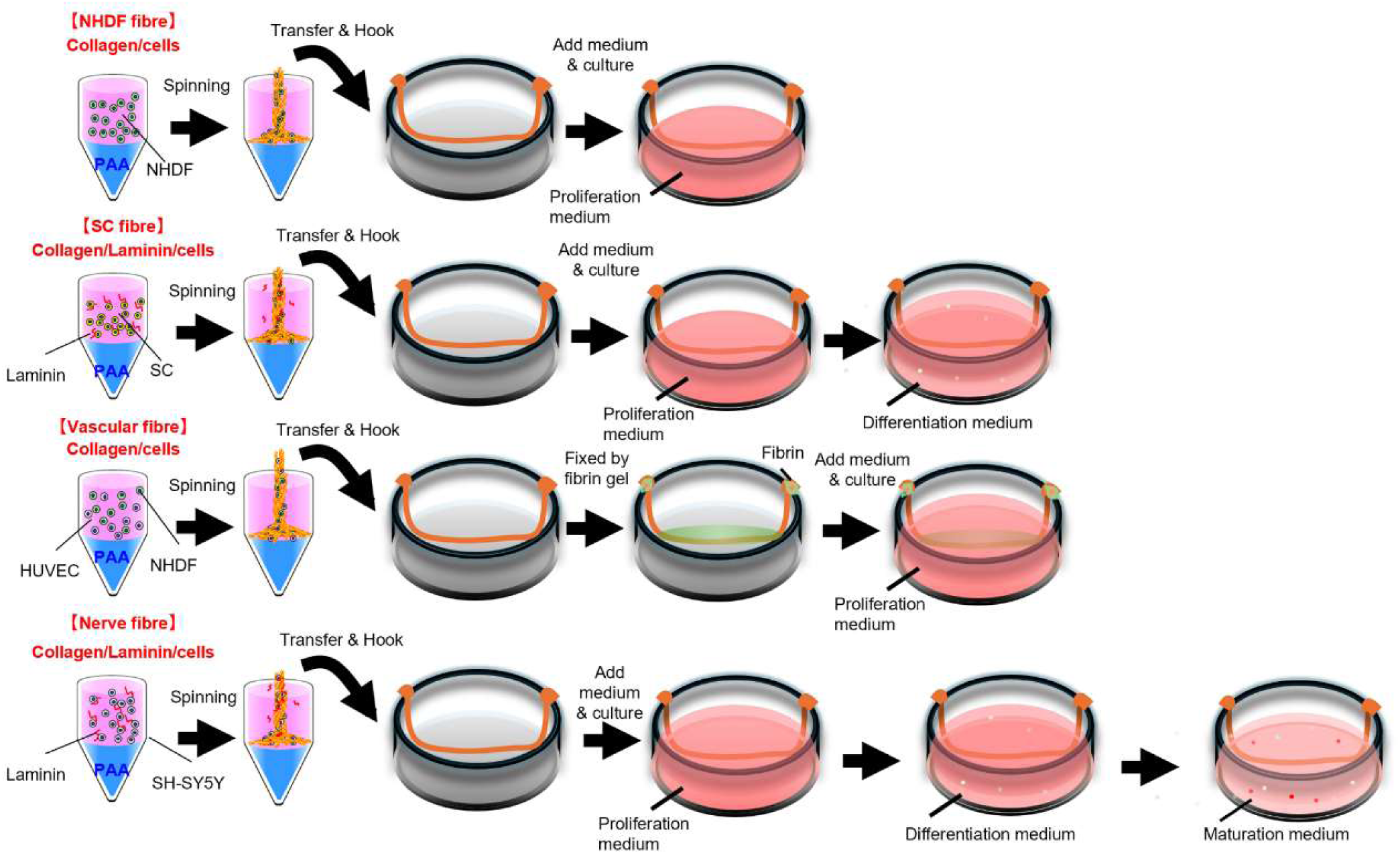
Schematic representation of fabrication and culture procedures for various cell-laden interfacial fibres. The illustration shows the composition of pre-spinning solutions and culture schedules for each tissue model: NHDF fibres, SC fibres, vascular fibres (NHDF–HUVEC co-culture), and nerve fibres (SH-SY5Y). Specific additives (e.g., laminin) were incorporated into the collagen solution depending on the target tissue. Culture phases—including proliferation, differentiation, and maturation—are indicated for each model. For vascular fibres, both ends of the spun construct were anchored with fibrin gel to prevent cell-mediated shrinkage during the proliferation phase.

**Extended Data Fig. 7.**
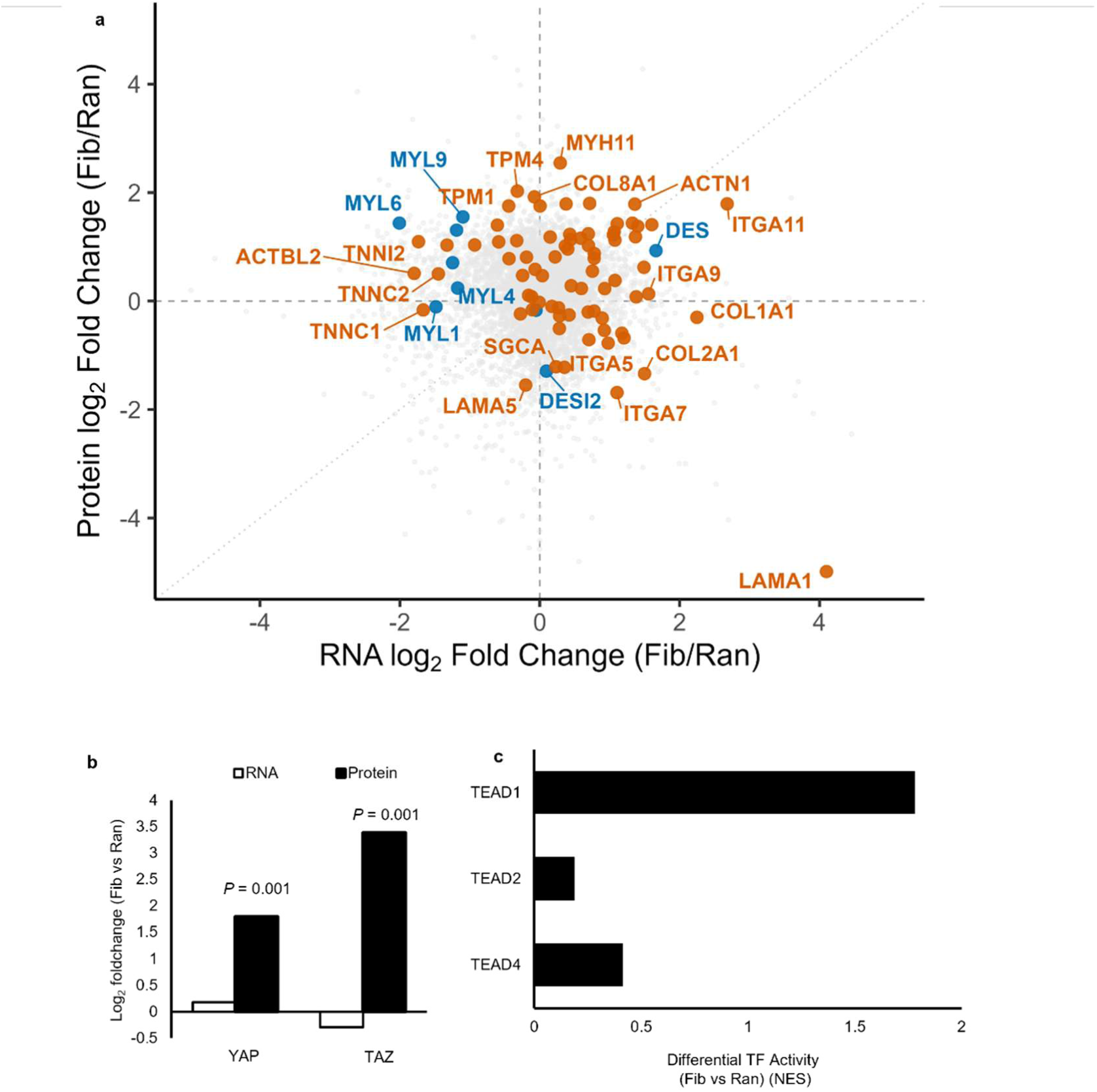
Transcriptomic and proteomic signatures underlying YAP/TAZ mechanotransduction. a, Transcriptome–proteome correlation plot (Fib vs. Ran). 2D scatter plot comparing log₂ fold changes of RNA (x-axis) and protein (y-axis) expression in aligned fibres (Fib) relative to random gel (Ran) (orange: cytoskeleton markers; blue: core sarcomere markers).**b,** Comparison of RNA and protein log₂ fold changes for mechanotransducers YAP and TAZ in Fib versus Ran.**c,** Inferred activity of TEAD family transcription factors. Bar plot showing differential transcription factor (TF) activity of TEAD1, TEAD2, and TEAD4 in Fib compared with Ran. TF activities were predicted from RNA-seq data, with the x-axis representing the difference in normalized enrichment scores (NES). Statistical significance was determined using an unpaired two-tailed Welch’s t-test.

**Extended Data Fig. 8.**
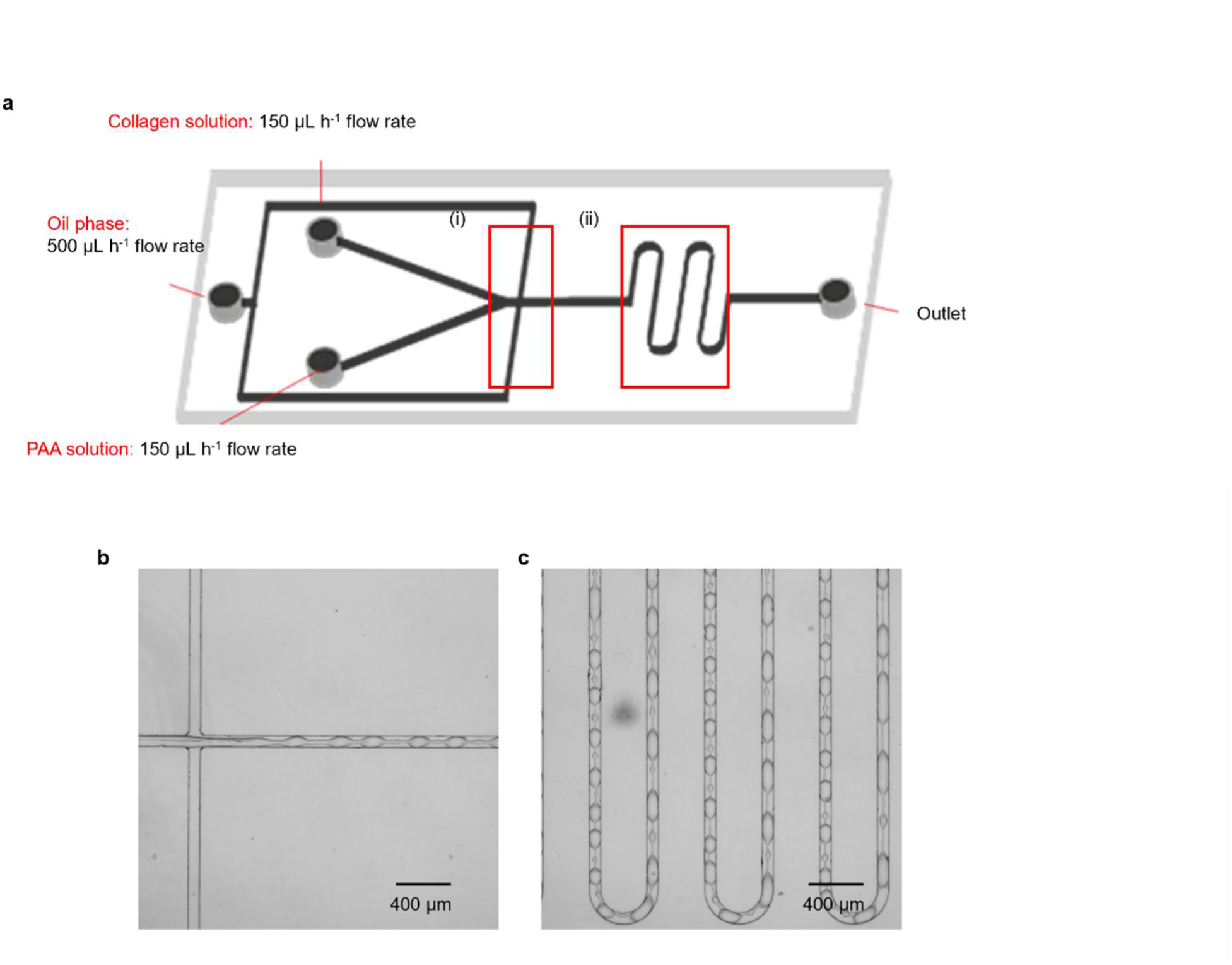
Microfluidic fabrication of necklace-like topological structures. a, Schematic illustration of the microfluidic device used for continuous generation of the interfacial complex. The red rectangles (i) and (ii) indicate the regions shown in b and c, respectively. Flow rates for the collagen solution, poly(acrylic acid) (PAA) solution, and the continuous oil phase were set to 150, 150, and 500 μL h⁻¹, respectively. **b,c,** Bright-field microscopy images showing the complex formation process at the channel junction (b) and the resulting necklace-like connected morphology in the downstream serpentine channel (c).

