## Supplementary Information for "Collagen Rope Trick: Cell-Laden Fibre Assembly at Liquid Interfaces"

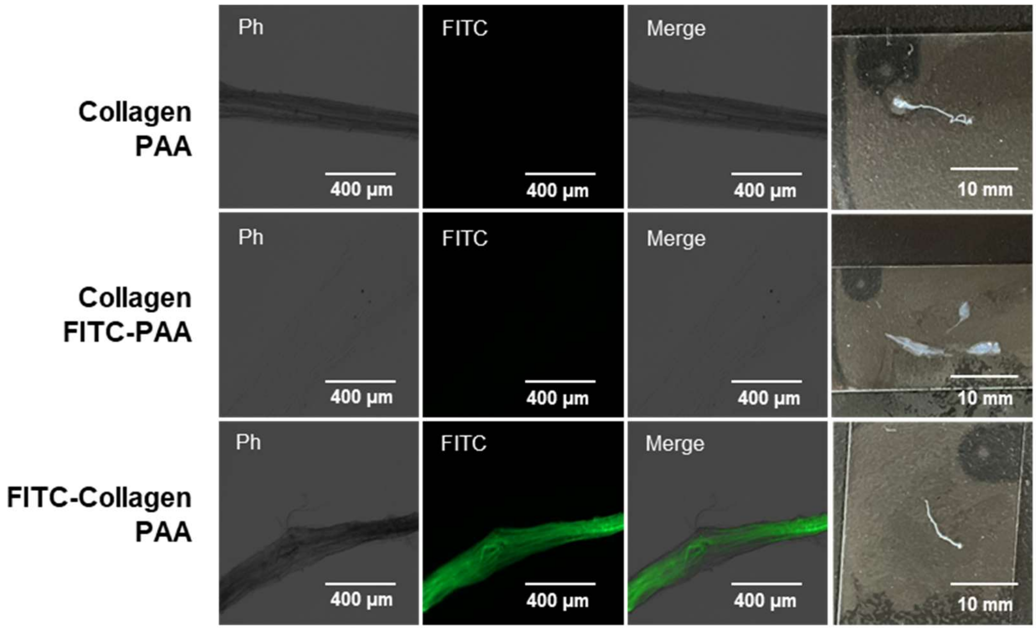


**Supplementary Fig. 1 | Fluorescence evaluation of the component polymers within the interfacial fibres.** Phase-contrast (Ph), fluorescence (FITC), and merged images of fibres fabricated using different combinations of labeled and unlabeled precursor solutions. Fibres were spun using unlabeled collagen (3 mg $mL^{-1}$), FITC-labeled collagen (1 mg $mL^{-1}$), unlabeled PAA (3 mg $mL^{-1}$ in PBS), or FITC-labeled PAA (3 mg $mL^{-1}$ in PBS). The images correspond to the unlabeled control (top), the FITC-PAA incorporated fibre (middle), and the FITC-collagen incorporated fibre (bottom). All fibres were thoroughly rinsed in ultrapure water before imaging.

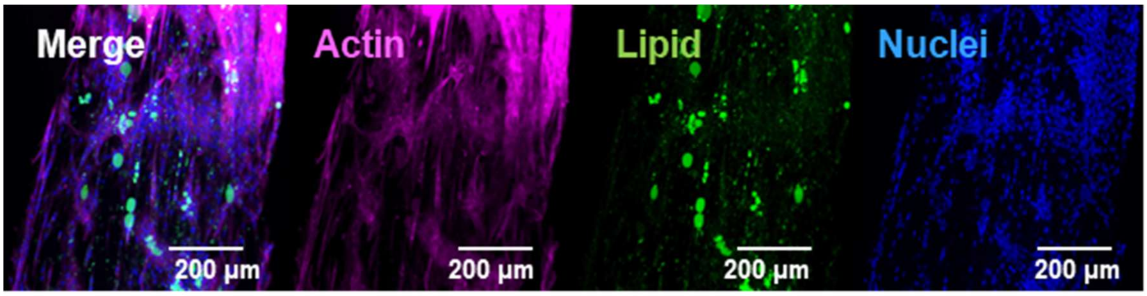


**Supplementary Fig. 2 | Immunofluorescence images of a bovine adipose fibre constructed using ADSCs.** The fibre is shown stained for specific markers (green: lipid droplets, magenta: actin, blue: nuclei). The fibre was cultured for 3 days for proliferation and 14 days for differentiation.

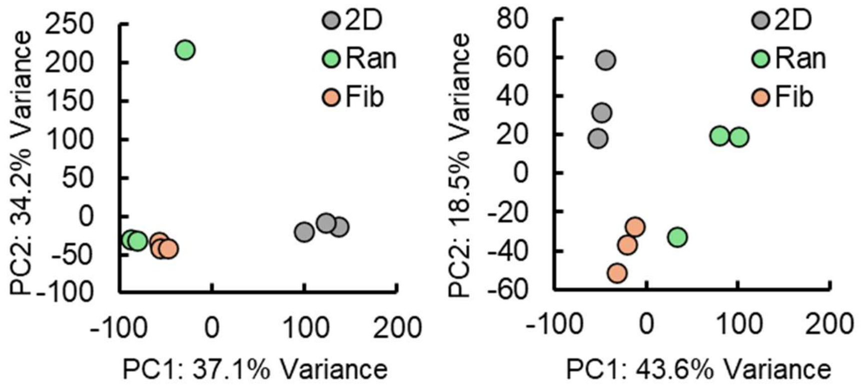


**Supplementary Fig. 3 | Principal component analysis (PCA) of transcriptomic and proteomic profiles.** PCA plots of RNA sequencing (transcriptome, left) and proteome (right) datasets across different culture conditions. Conditions include traditional 2D culture (2D, gray circles), random 3D collagen constructs (Ran, green circles), and fabricated aligned collagen fibres (Fib, orange circles). Percentages on the axes indicate the proportion of total variance explained by principal component 1 (PC1) and principal component 2 (PC2).

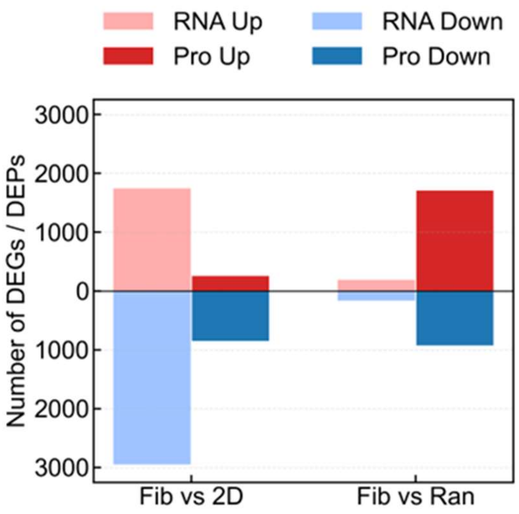


**Supplementary Fig. 4 | Quantification of differentially expressed genes (DEGs) and proteins (DEPs).** Total number of significantly up-regulated (positive y-axis) and down-regulated (negative y-axis) features in aligned fibres (Fib) compared with traditional 2D culture (Fib versus 2D) and random 3D constructs (Fib versus Ran). Light red and light blue bars represent up- and down-regulated RNAs (DEGs), respectively. Dark red and dark blue bars represent up- and down-regulated proteins (DEPs), respectively. Significant DEGs and DEPs were defined by an adjusted *P* value (false discovery rate, FDR) $< 0.05$ and an absolute $\log_2$ fold change $\geq 0.585$.

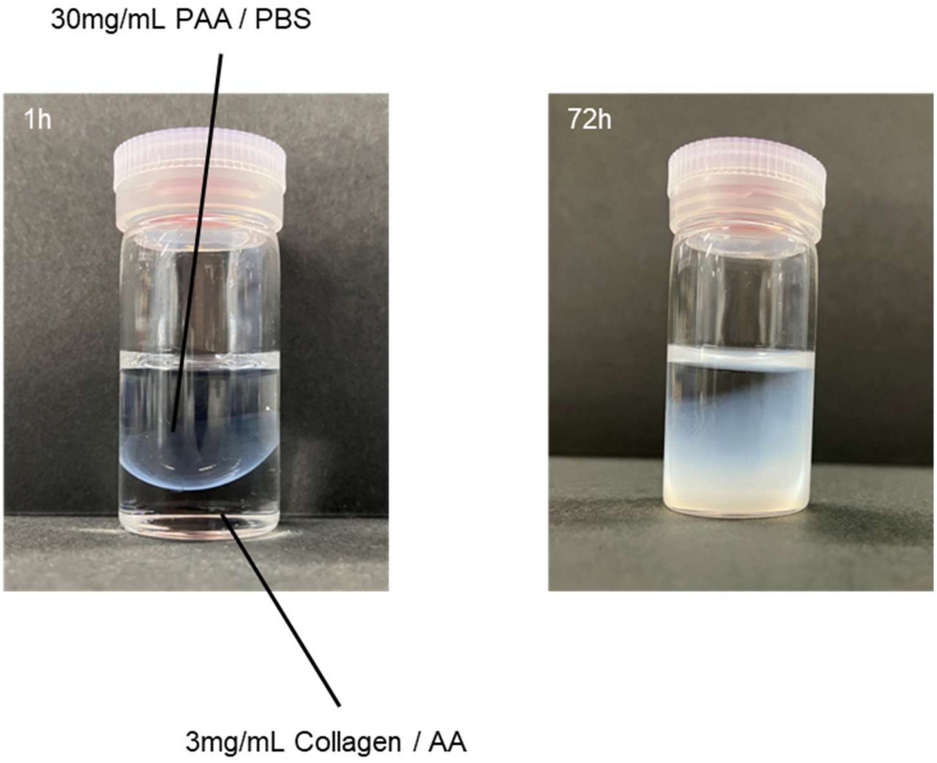


**Supplementary Fig. 5 | Time-dependent formation and growth of the collagen–PAA interfacial sheet.** Macroscopic appearance of the interface development at 1 h (left) and 72 h (right). A PAA solution (30 mg $mL^{-1}$ in PBS) was gently layered onto a collagen solution (3 mg $mL^{-1}$ in acetic acid) to create a liquid–liquid interface. This procedure corresponds to the fabrication process for the sheets presented in **Fig. 5a**.

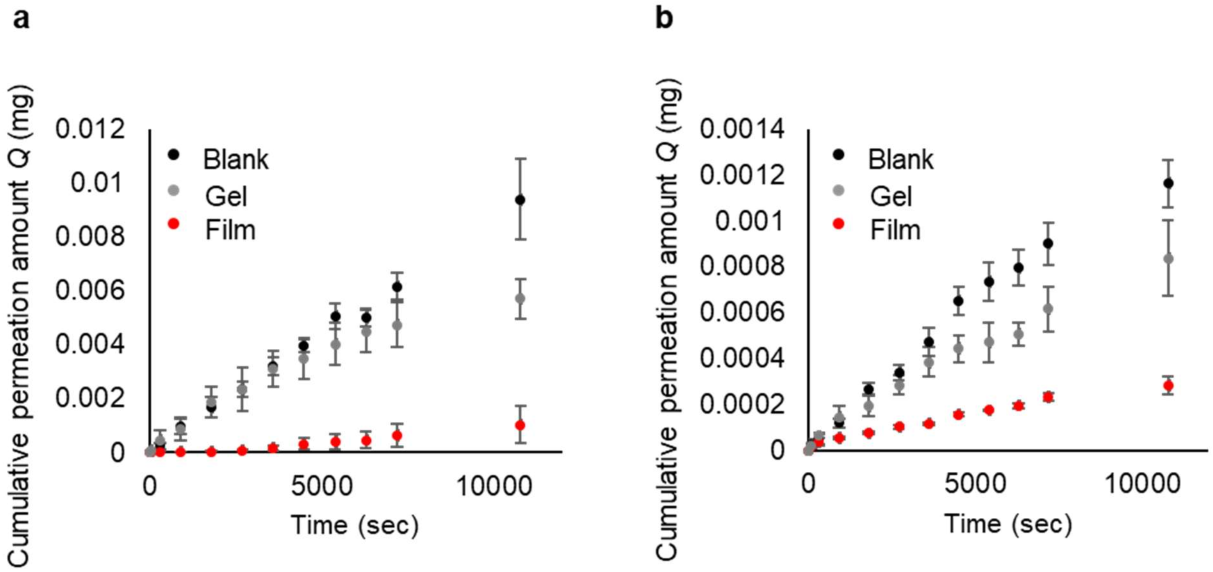


**Supplementary Fig. 6 | Permeation kinetics of fluorescent dextrans. a-b,** Time-dependent cumulative permeation amount (*Q*) of 4 kDa FITC-dextran (**a**) and 2,000 kDa TRITC-dextran (**b**). Permeation profiles are shown for the fabricated interfacial collagen film (red circles), bulk collagen gel (gray circles), and the cell-free insert membrane (black circles). The steady-state flux, corresponding to the slope of the linear region of these curves, was used to calculate the apparent permeability ($P_{app}$) and the effective permeability coefficient ($P_e$) presented in **Fig. 5b.** Error bars represent ± s.d. (*n* = 3 independent samples)

| Reference | Category | Material | Fracture strength (MPa) | Toughness (MJ m$^{-3}$) |
|---|---|---|---|---|
| 1. Wu, J. et al. *Adv. Funct. Mater.* (2023), **33**, 2210395 | Supramolecular hydrogel | Poly (NASC-co-AA) | 9.1 | 33.7 |
| 2. Hu, X. et al., *Adv. Mater.* (2015), **27**, 6831-6837 | Hydrogen-bonded hydrogel | DMAA-co-MAAc | 28 | 80 |
| 3. Wang, M. et al., *Nature* (2024), **631**, 313–318 | Traditional hydrogel | PAA-H2O | 0.3 | 0.2 |
| 4. Sun, T. L. et al., *Nat. Mater.* (2013), **12**, 932–937 | Polyampholyte hydrogel | Poly(NaSS-co-MPTC) | 2 | 7 |
| 5. Sato, K. et al., *Adv. Mater.* (2015), **27**, 6990–6998 | Phase-separated hydrogel | PAAm | 6 | 50 |
| 6. Li, L. et al., *Angew. Chem. Int. Ed.* (2022), 61, e202212512 | Tough ionogel | PDMAA / [Bmim][ZnX] | 14.3 | 78 |
| 7. Gong, J. P., *Soft Matter* (2010), 6, 2583–2590 | B-DN hydrogel | PAMPS/PAAm | 17 | 10 |
| 8. Gallagher, A. J. et al., *IRCOBI Conf.* (2012), **2012**,494-502 | Skin | Human back skin | 27.2 | 4.9 |
| 9. Pita, V. J. R. R. et al., *Polym. Test.* (2002), **21**, 545–550 | Plasticized PVC | PVC / DIDP | 10 | 15.7 |
| 10. Yarger, J. L. et al., *Nat. Rev. Mater.* (2018), **3**, 18008 | Tendon collagen | Mammalian tendon | 120 | 6 |
| 11. Yarger, J. L. et al., *Nat. Rev. Mater.* (2018), **3**, 18008 | Spider dragline silk | Spider silk | 350 | 140 |
| 12. Corkhill, P. H. et al., *Proc. Inst. Mech. Eng. H.* (1990) 204, 147–155 | Cartilage | Cartilage | 20 | 30 |
| This work | This work | Collagen | 280 | 17 |

**Supplementary Table 1 | Comparison of mechanical properties between the collagen fibre in this work and various materials reported in the literature[1–11] in reference section.** NASC, *N*-acryloylsemicarbazide; AA, acrylic acid; DMAA, *N,N*-dimethylacrylamide; MAAc, methacrylic acid; PAA, poly(acrylic acid); NaSS, sodium *p*-styrenesulfonate; MPTC, 3-(methacryloylamino)propyl-trimethylammonium chloride; PAAm, polyacrylamide; PDMAA, poly(*N,N*-dimethylacrylamide); [Bmim][ZnX], 1-butyl-3-methylimidazolium halozincate; PAMPS, poly(2-acrylamido-2-methylpropanesulfonic acid); PVC, poly(vinyl chloride); DIDP, di-isodecyl phthalate. The maximum reported values for tensile stress and toughness (work of extension) from each reference are summarized.

**Supplementary Video 1** Phase-contrast microscopy video of the horizontal drawing of a collagen fiber at the liquid-liquid interface. Poly(acrylic acid) (PAA) solution is initially applied on the left, followed by the addition of the collagen solution to establish contact. The video demonstrates the continuous spinning of a collagen fiber by horizontally pulling the newly formed solid-like interface with forceps.

**Supplementary Video 2** Phase-contrast microscopy video of instantaneous interface formation. A collagen solution is dropped into a poly(acrylic acid) (PAA) solution, resulting in the immediate formation of a distinct boundary. The video captures the solid-like interfacial membrane progressively thickening towards the collagen phase over time.

**Supplementary Video 3** Macroscopic video of the continuous interfacial spinning process. A poly(acrylic acid) (PAA) solution, stained with blue food coloring for visualization, is dropped into a collagen solution prepared in a 35 mm dish. The video demonstrates the continuous generation of a macroscopic fiber by vertically drawing the formed interface with forceps.

**Supplementary Video 4** High-speed macroscopic video of the continuous interfacial spinning process. A collagen solution is layered on top of a poly(acrylic acid) (PAA) solution. The video, captured at 1,000 frames per second (fps) and replayed at 120 fps, demonstrates the continuous vertical drawing of the interfacial membrane using a fine pipette tip.

**Supplementary Video 5** Phase-contrast microscopy video of the horizontal drawing of a cell-laden collagen fiber. A cell-suspended collagen solution is initially applied, followed by the addition of a poly(acrylic acid) (PAA) solution to establish contact. The video demonstrates the continuous spinning process by horizontally pulling the newly formed interface with forceps, illustrating how the suspended cells are actively incorporated into the forming fiber.

**Supplementary Video 6** 3D reconstructed video of an SC-laden fiber acquired by light-sheet fluorescence microscopy. The fiber is immunostained for myosin heavy chain (magenta). The video provides a volumetric view of the construct, demonstrating the formation of highly aligned and differentiated myotubes deep within the interior of the fiber.

**Supplementary Video 7** 3D reconstructed video of an adipose-derived stem cell (ADSC) capsule. The construct is immunostained for Col I (magenta) and actin (green). The video provides a volumetric view of the collagen capsule, demonstrating that the cells are uniformly attached and well-spread throughout the entire surface and interior of the collagen matrix.
